# A novel CCDC112-dependent process of sperm midpiece formation and epididymal maturation

**DOI:** 10.1101/2024.11.05.622046

**Authors:** Maddison L. Graffeo, Joseph Nguyen, Farin Yazdan Parast, D. Jo Merriner, Jessica E.M. Dunleavy, Denis Korneev, Hidenobu Okuda, Anne E. O’Connor, Donald F. Conrad, Reza Nosrati, Brendan Houston, Moira K. O’Bryan

## Abstract

The sperm mitochondrial sheath has proposed functions in structural support and energy production for motility. Here we define coiled coil domain containing protein 112, CCDC112, as crucial for male fertility, specifically in the assembly and function of the mitochondrial sheath. We unveiled a previously unrecognised process of epididymal mitochondrial sheath maturation. Sperm mitochondrial sheaths are structurally immature upon exiting the testis and continue to mature as they transit from the caput to the cauda epididymis. Data reveal the critical role of CCDC112 in mitochondrial morphogenesis during sheath formation. A lack of CCDC112 lead to significantly reduced mitochondrial respiration capacity, irregular flagellar waveforms, diminished progressive motility and a failure of sperm to traverse the female reproductive tract to the site of fertilisation and penetrate the zonae pellucidae of oocytes. Finally, we identify CCDC112 as a component of the distal appendages of the mother centriole and identify a facilitative role in core tail assembly. Collectively, CCDC112 is a key regulator of sperm mitochondrial sheath assembly while further expanding the understanding of spermatogenesis, energy generation and flagella kinematics.

## Introduction

Mammalian sperm are streamlined, highly polar cells, optimised for traversing the female reproductive tract, locating and fertilising an ovum. The sperm flagellum is essential to this process (1). It is composed of a central microtubule-based axoneme, which functions as the ‘drive shaft’ of the tail, surrounded by three region-specific accessory structures: the outer dense fibres (mid– and principal pieces), the mitochondrial sheath or midpiece (midpiece specific) and the fibrous sheath (principal piece specific). The processes underpinning sperm assembly remain poorly defined at a molecular level but involve a series of complex microtubule-based protein and organelle transport systems (2). The mitochondrial sheath, is hypothesized as essential for both structural support (3, 4) and as a source of ATP, via oxidative phosphorylation, to fuel axoneme movement and thus sperm motility (5, 6). The precise role of mitochondrial generated ATP in sperm function, however, remains controversial (5, 6). Such controversy arises in part due to differences in assay conditions between studies (7) and likely genuine differences in sperm metabolism between species (5, 8–15).

In addition to mitochondrial ATP generation in the midpiece, sperm also produce ATP by glycolysis in the principal piece (16–18). In somatic cells, glycolysis and oxidative phosphorylation are usually tightly coupled to maximise ATP production (19). Conversely, in sperm, the majority of glycolytic enzymes are anchored to fibrous sheath in the principal piece, and to a lesser extent in the sperm head and midpiece (16–18, 20, 21), whereas oxidative phosphorylation occurs solely in the midpiece. As such, coupling between the two systems is not obvious with the traditional dogma suggesting they function separately. Recent data, however, challenges this notion, proposing crosstalk between the two systems (22–24).

The formation of the mitochondrial sheath at an organelle level has been described, initiating via the recruitment, migration, and successive alignment of spherical mitochondria from the cytoplasmic lobes of elongated spermatids toward the axoneme (25, 26). Mitochondria then attach to the outer dense fibres surrounding the axoneme along the portion that will ultimately become the midpiece. They then undergo dramatic morphological modifications, transforming from individual spherical organelles into a double helix of rod-shaped mitochondria that abut end to end in the compacted sheath (25, 26). The process by which the mitochondrial sheath is assembled at a molecular level, and the importance of the structure itself to male fertility, however, remain poorly defined. What is clear is that abnormalities in mitochondrial sheath structure are associated with mammalian male infertility including in humans (27–29) and mice (27, 30–34). Equally, evolutionary studies in humans and mice have shown that sperm midpiece length is positively associated with sperm swimming speed (35, 36). Mitochondrial volume is also significantly correlated with swimming speed, in addition to, flagellar beat frequency and ATP content (35, 37) whereas outer dense fibre volume is only positively associated with ATP content (35).

The coiled coil domain containing (CCDC) protein superfamily is a large group of diverse proteins characterised by their coiled coil domains, which are comprised of two or more entwined alpha helices (38). They function by facilitating protein-protein interactions, potentially acting as molecular spacers by scaffolding macromolecular complexes or separating functional protein domains in a wide range of biological contexts during cilia and flagella biogenesis, including sperm tails (39–44), centrosome organisation (39, 45, 46) and fertilisation (44). One such member, CCDC112, has recently been proposed as a ciliation factor in cilia producing RPE-1 somatic cells, colocalising with PCM1 in the pericentriolar material (47, 48). Depletion of CCDC112 in RPE-1 cells results in perturbed centriolar satellite intensity (47). Single cell sequencing data revealed that CCDC112 is highly expressed in pachytene and diplotene spermatocytes in human testes (49).

Herein, we test the function of CCDC112 *in vivo,* demonstrating an essential role for CCDC112 in male fertility, specifically in mitochondrial morphogenesis and remodelling during mitochondrial sheath formation. Loss of CCDC112 function results in male sterility due to a highly abnormal mitochondrial sheath structure, reduced ATP production, and an irregular flagella waveform with insufficient mechanical power to drive progressive sperm motility and fertilisation. We further identify CCDC112 as a distal appendage component of the mother centriole, thus identifying a potential mechanism for the development of a sub-population of short sperm in the absence of CCDC112. Finally, we reveal a previously unrecognised form of epididymal sperm maturation, epididymal mitochondrial sheath maturation. Collectively, these data identify CCDC112 as a critical regulator of sperm midpiece assembly and function and uncover a novel process required for male fertility.

## Results

### CCDC112 is predominantly expressed in the testis

*CCDC112* is a highly conserved gene between human and mouse with 93% protein similarity. A previously published analysis of *CCDC112* expression in human organs revealed it was highly enriched within testes compared to other tissues (*p* < 0.0001; Fig. S1A) (50). Further single cell RNA sequencing data of germ cells revealed that in humans *CCDC112* is expressed most highly expressed in late primary spermatocytes undergoing the process of meiosis and to a lesser extent in spermatogonia (Fig. S1B) (51). A similar pattern of expression was observed in mouse male germ cells (Fig. S1C) (52). Unfortunately, and despite assessing four CCDC112 antibodies, including one produced in-house, all antibodies tested in this study bound non-specifically to other proteins in immunochemistry and western blotting protocols. As such, we were unable to define the localisation of CCDC112.

### CCDC112 is essential for male fertility

To explore the role of CCDC112 in male fertility, we produced a *Ccdc112* knockout mouse model (*Ccdc112^KO/KO^*) whereby we removed exon 2 from the principal *Ccdc112* mouse transcript (*ENSMUST00000072835.7)*, which introduced a premature stop codon in exon 3 (Fig. S1E, F). Quantitative PCR measured a 99.8% reduction in *Ccdc112* mRNA expression in testis compared to wildtype littermates thus confirming exon removal in *Ccdc112^KO/KO^* animals (Fig. S1D).

*Ccdc112^KO/KO^* mice appeared outwardly normal with no overt behavioural abnormalities and displayed a normal mating frequency. They were, however, male sterile (0 pups per copulatory plug versus 7.5 in *Ccdc112^WT/WT^*, p < 0.0001; Fig. 1A). Body weights (Fig. 1B) were comparable between *Ccdc112^WT/WT^* and *Ccdc112^KO/KO^*mice. While testis weights were unchanged between genotypes (Fig. 1C), testis daily sperm production (DSP) and epididymal sperm content (ESC), were significantly reduced by 13% (p < 0.05) and 38% (p < 0.001), respectively, in *Ccdc112^KO/KO^* males compared to *Ccdc112^WT/WT^* littermates (Fig. 1D, E). This divergence between testis weight and sperm output is suggestive of a loss of spermatids during late spermiogenesis, wherein they contributed minimally to overall testis weight. Further, the more severe reduction in ESC compared to DSP is consistent with a partial spermiation failure illustrated by the presence of retained sperm in stage IX *Ccdc112^KO/KO^*tubules (Fig. 1H), but not *Ccdc112^WT/WT^* mice. Additionally, while both the number of round spermatids in stage VIII tubules (Fig. S2A) and apoptotic germ cells across all tubule stages (Fig. S2B) were comparable between genotypes, we observed elevated levels of sloughed round germ cells in the *Ccdc112^KO/KO^*epididymides. Germ cell sloughing was rarely observed in *Ccdc112^WT/WT^*mice (Fig. 1I).

**Figure 1.**
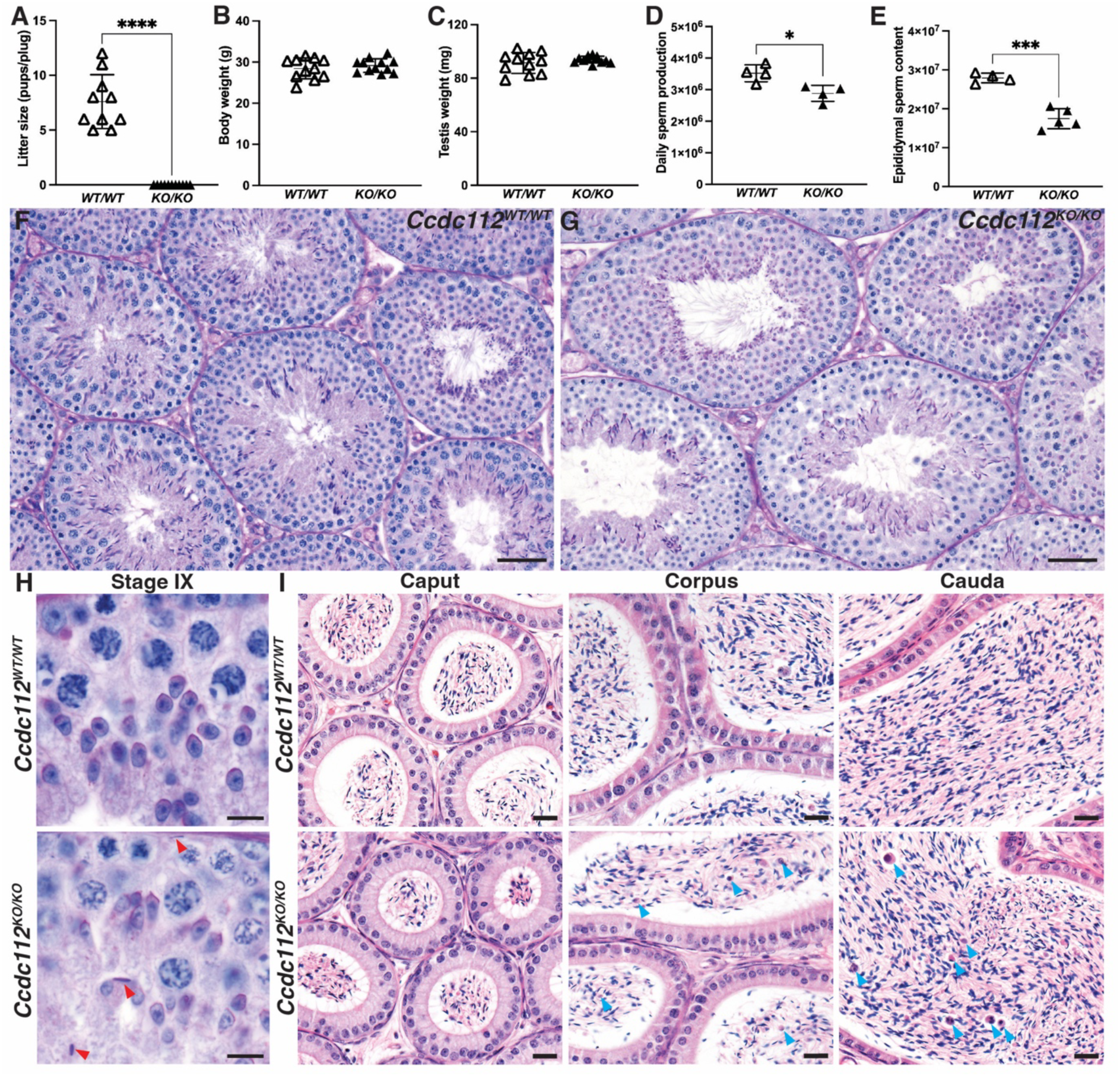
CCDC112 is required for male fertility. Litter size (A), body weight (B), testis weight (C), testis daily sperm production (DSP) (D) and total epididymal sperm count (ESC) (E) in *Ccdc112^WT/WT^* mice and *Ccdc112^KO/KO^*mice (n ζ 4/genotype). Periodic acid-Schiff stained testis sections (F-H) and haematoxylin and eosin-stained epididymis sections (I) from *Ccdc112^WT/WT^* and *Ccdc112^KO/KO^*mice. Red and blue arrowheads indicate retained spermatids in (H) and prematurely sloughed spermatids in (I), respectively. Lines denote mean ± SD in A-E; * P <0.05, *** P <0.001, **** P <0.0001. Scale bars in F-G = 50 μm, H = 10 µm, I = 20 µm.

Cauda epididymal sperm from *Ccdc112^KO/KO^* males exhibited a significantly reduced ability to achieve any form of motility (85% in *Ccdc112^WT/WT^* versus 52.4% in *Ccdc112^KO/KO^*; p < 0.01, Fig. 2A) and notably progressive motility (50.2% in *Ccdc112^WT/WT^* versus 15.6% in *Ccdc112^KO/KO^*; p < 0.001, Fig. 2B). Those sperm that did display progressive motility swam on average more slowly than sperm from wildtype males (76.6 µm/s in *Ccdc112^WT/WT^* versus 56.7 µm/s in *Ccdc112^KO/KO^*; p < 0.0001, Fig. 2C).

**Figure 2.**
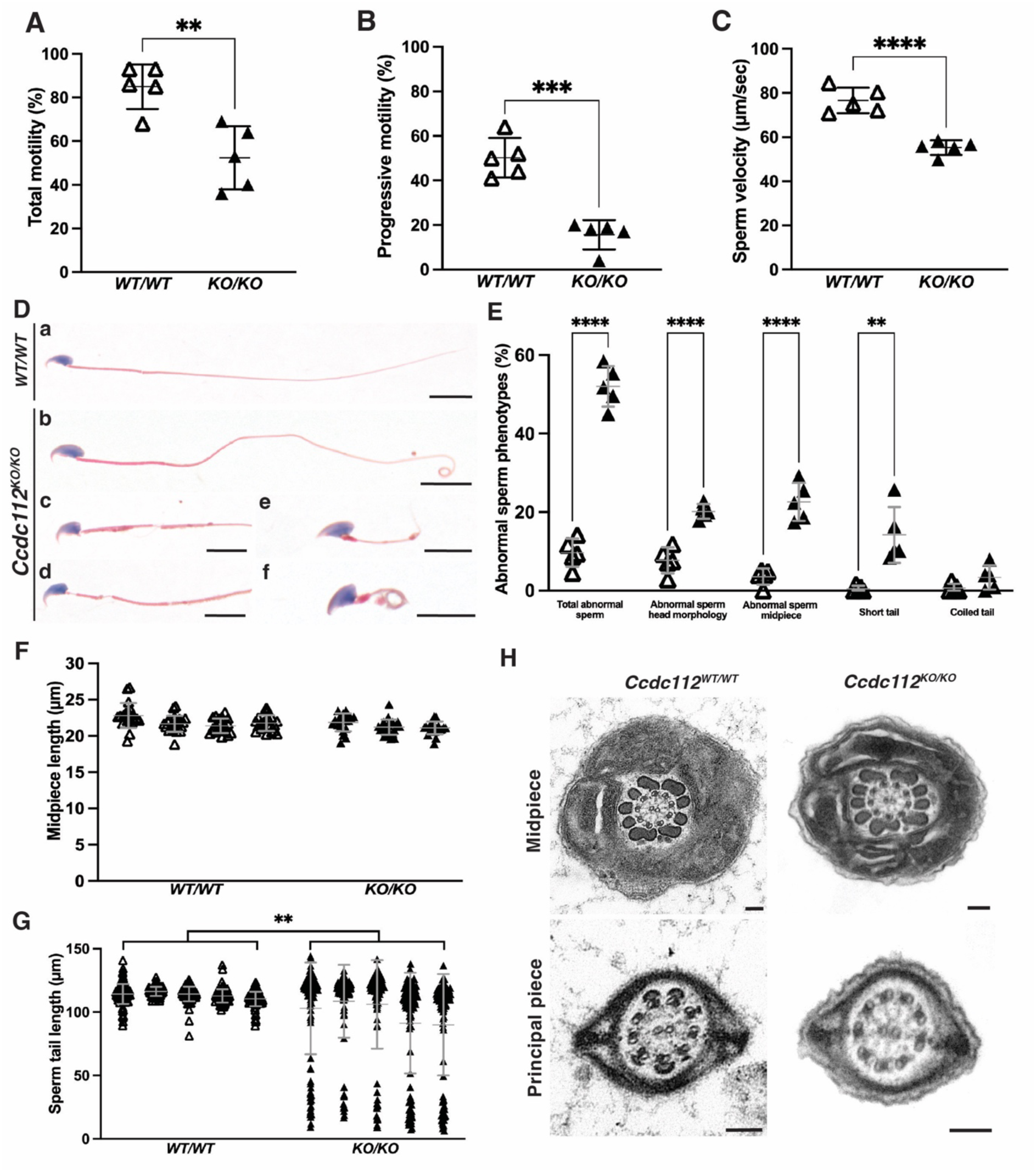
CCDC112 is essential for sperm tail structure and function. Percentage of motile (A) and progressively motile sperm (B) and average sperm velocity (C) of cauda epididymal sperm from *Ccdc112^WT/WT^* mice and *Ccdc112^KO/KO^* mice (n = 5 mice/genotype). (D) Cauda epididymal sperm morphology as stained by haematoxylin and eosin. Abnormalities were observed in a majority of *Ccdc112^KO/KO^* sperm, including abnormal sperm head morphology (c-f), mitochondrial sheath defects (c, d), short (e) and coiled (f) sperm tails. (E) Quantification of abnormal sperm morphology phenotypes in *Ccdc112^WT/WT^*(white triangles) and *Ccdc112^KO/KO^* males (black triangles) (n = 5/genotype). Similarly, on these sperm, midpiece compartment (F) and total tail (G) length were measured (n = 3-5/genotype). (H) Axoneme ultrastructure of *Ccdc112^WT/WT^*and *Ccdc112^KO/KO^* cauda epididymal sperm as assessed via transmission electron microscopy. Lines denote mean ± SD in A-C, E-G; ** P <0.01, *** P <0.001, **** P < 0.0001. Scale bars in D = 10 μm, H = 100 nm.

An examination of *Ccdc112^KO/KO^* cauda epididymal sperm structure at a light microscopic level revealed that the majority were morphologically abnormal (52.02% abnormal in *Ccdc112^KO/KO^*versus 9.66% in *Ccdc112^WT/WT^*; p < 0.0001, Fig. 2D, E). Specifically, intact sperm from *Ccdc112^KO/KO^* males presented with abnormal sperm head morphology (20.2% versus 7.7% *Ccdc112^WT/WT^*; p < 0.0001, Fig. 2D, E), obvious defects in midpiece structure (22.6% versus 3.6%; p < 0.0001, Fig. 2Dc, d, E) and a higher incidence of short sperm tails (14.2% versus 0.5%; p < 0.01, Fig. 2De, E). While sperm produced by *Ccdc112^KO/KO^* males had normal midpiece (mitochondrial sheath) lengths (Fig. 2F), a significant reduction was found in average principal piece and total tail length (p = 0.0089, Fig. 2G). Sperm fell into two major clusters – those where no significant differences in total sperm tail length and those where the tail was significantly shorter (p < 0.0001). An assessment of the sperm axoneme structure via transmission electron microscopy revealed no difference in structure (Fig. 2H). In the majority of sperm, accessory structures including the outer dense fibres and fibrous sheath were present and superficially normal when viewed in a transverse plane. Further analysis of sperm head morphology defects via testis section staining for α-tubulin as a marker of the manchette, a microtubule skirt-like structure essential for sperm head shaping, revealed comparable assembly, migration and disassembly of the manchette between genotypes (Fig. S3). Additional testing to determine the origin of the head morphology defects is required.

Given that 15.6% of sperm in *Ccdc112^KO/KO^* exhibit progressive motility, fertilisation remained possible. To test why this did not occur *in vivo*, we assessed the ability of sperm to ascend the female reproductive tract following natural mating (Fig. 3A). Compared to wildtype, 66% fewer sperm from *Ccdc112^KO/KO^* males were present in the isthmus of the Fallopian tube (p = 0.01) and 87% fewer in the ampulla (p = 0.04). To extend this analysis, we examined the consequences of CCDC112 loss on sperm function in *in vitro* fertilisation wherein the majority of physical barriers to fertilization were removed. Sperm from wildtype males achieved efficient fertilisation, leading to a two-cell development rate of 90.62% (Fig. 3B). By contrast, only 12.09% of oocytes were fertilised by sperm from *Ccdc112^KO/KO^* males (p < 0.0001, Fig. 3B). A visual inspection revealed that the loss of CCDC112 impaired the ability of sperm to penetrate the zonae pellucidae (Fig. S4).

**Figure 3.**
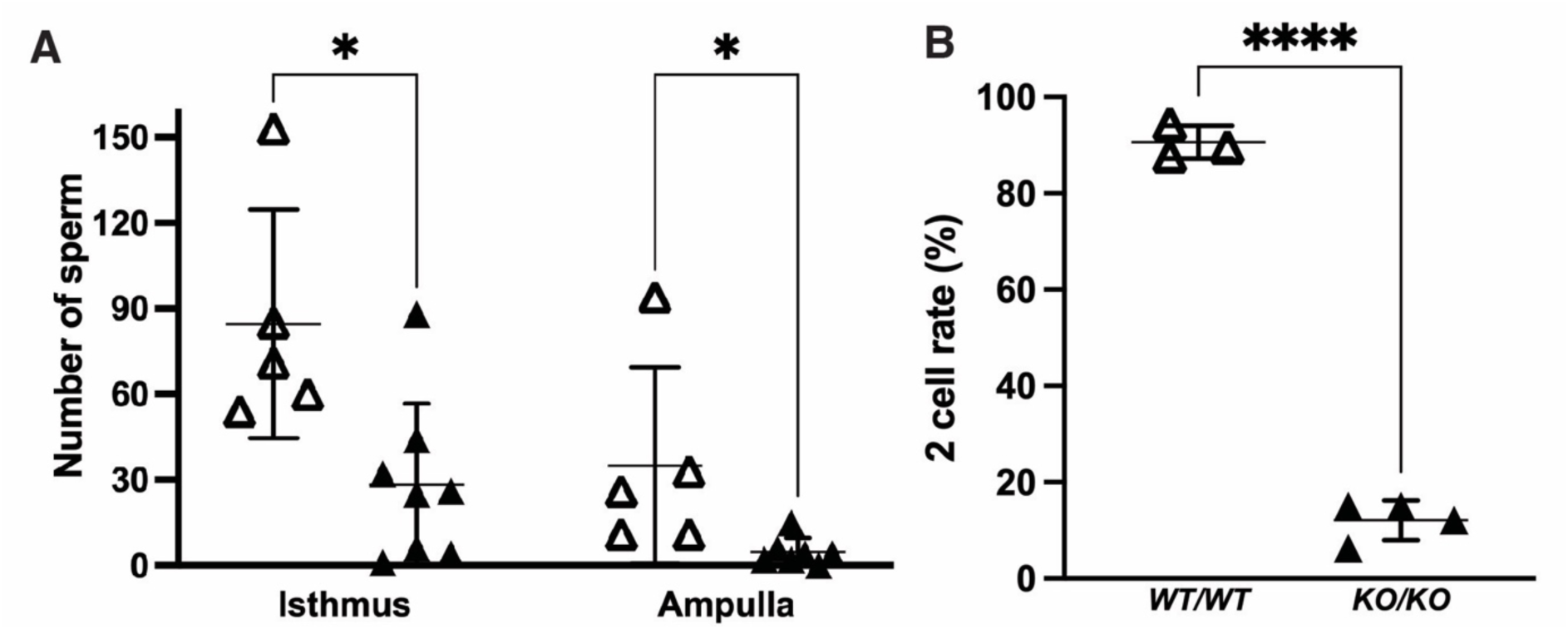
CCDC112 is essential for sperm transit through the female reproductive tract and fertilisation. (A) Quantification of *Ccdc112^WT/WT^* (white triangles) and *Ccdc112^KO/KO^* mouse sperm (black triangles) in the isthmus and ampulla region of the oviduct in the female reproductive tract (n ≥ 5/genotype). (B) Percentage of oocytes which developed to the two-cell stage after fertilisation with *Ccdc112^WT/WT^* and *Ccdc112^KO/KO^*mouse sperm (n ≥ 3/genotype).

CCDC112 is thus an essential requirement for mouse sperm to reach the site of fertilisation in the female reproductive tract and to penetrate the outer vestments of the oocyte.

### CCDC112 regulates mitochondrial alignment and organisation in the sperm midpiece

To explore the consequences of CCDC112 loss on sperm morphology more precisely, we used scanning electron microscopy on plasma membrane stripped sperm from the wildtype mouse caput and cauda epididymides (Fig. 4A). We examined the organisation of the mitochondrial and fibrous sheaths in wildtype sperm. The results revealed a previously unappreciated finding – that the quality of sperm mitochondrial sheath packing in wildtype sperm improves between the caput (proximal region) and the cauda (distal region) epididymis (Figure 4), suggestive of a novel form of epididymal sperm maturation. A qualitative overview of sperm from the caput epididymis revealed that the majority of sperm contained mitochondrial defects, including misaligned, incorrectly oriented, and morphologically abnormal (inconsistent shape and size) mitochondria. By contrast, for wildtype mouse cauda epididymal sperm the majority of the most severely abnormal midpiece phenotypes, including immature (spherical and/or crescent shaped) mitochondria, absent mitochondria, and/or exposed outer dense fibres were no longer present (Fig. 4B, F). A similar but more severe range of defects were observed in sperm from *Ccdc112^KO/KO^* males. Additional defects included the complete absence of mitochondria and large gaps in the midpiece (Fig. 4F).

**Figure 4.**
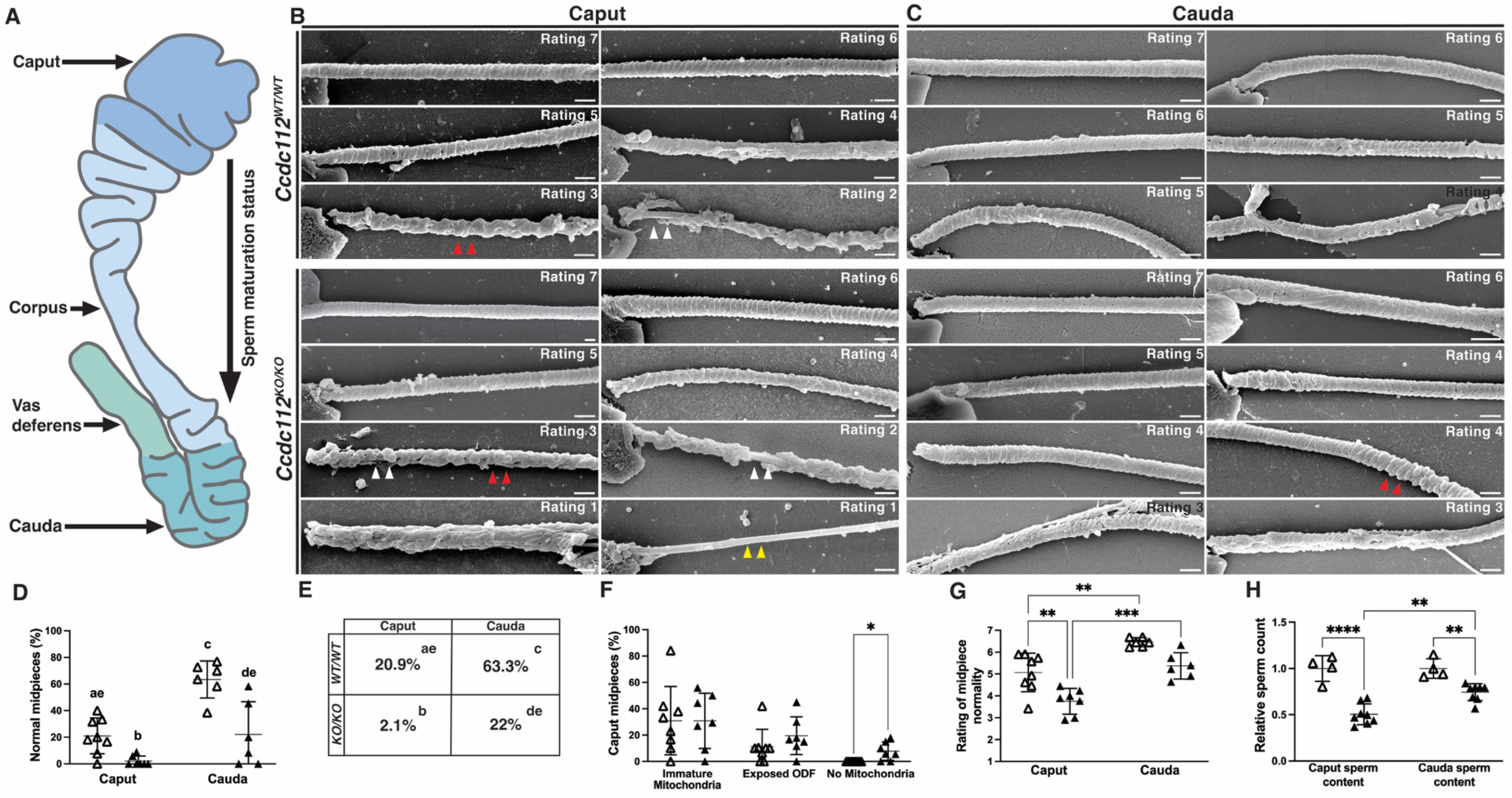
CCDC112 regulates mitochondrial sheath formation. (A) A schematic of the epididymis in mice. Once sperm are released from the testis they are transported to the epididymis for maturation, quality control, and storage. They first enter into the caput (head) epididymis before transiting through the corpus (body) towards the cauda (tail) before eventually exiting through the vas deferens upon ejaculation. Scanning electron micrography images of caput (B) and cauda (C) epididymal sperm midpieces from *Ccdc112^WT/WT^*and *Ccdc112^KO/KO^* mice. Midpieces were rated on a scale of 1 to 7, based on mitochondrial sheath abnormality severity. A score of 1 represents high disorganised and abnormal mitochondrial sheath formation and a score of 7 represents highly organised mitochondrial alignment, i.e., normal mitochondrial sheath formation. No rating 1 sperm were seen in *Ccdc112^WT/WT^* samples. Abnormalities included irregular alignment, morphology and size of mitochondria, immature mitochondria (red arrowheads), exposed outer dense fibres (white arrowheads; ODFs) and no mitochondria (yellow arrowheads). Percentage of normal caput and cauda midpieces (D-E) in *Ccdc112^WT/WT^*(white triangles) and *Ccdc112^KO/KO^*(black triangles) males (n ζ 10 midpieces/animal and n ζ 4/genotype). Differing lowercase letter designations denote significant differences between groups in D-E. Percentage of caput midpieces with immature ODF and/or no mitochondria (F), the rating of caput and cauda epididymal sperm midpiece normality (G), and the relative difference between caput sperm content and cauda sperm content (H) in *Ccdc112^WT/WT^* and *Ccdc112^KO/KO^*males (n ζ 10 midpieces/animal and n ζ 4/genotype). Lines denote mean ± SD in D, F-H; * P < 0.05, ** P < 0.01, *** P < 0.001, **** P < 0.0001. Scale bars in B-C = 0.5 μm.

To quantify these defects, sperm midpiece structure was scored on a normality scale (1–7) between epididymal regions and genotypes. In wildtype caput epididymides, 20.9% of sperm present were classified as high quality (normal, rating 7) compared to 63.3% of sperm harvested from the cauda region, thus highlighting the improvement in sperm structural quality that occurs during epididymal transit (p < 0.0001, Fig. 4D-E). For *Ccdc112^KO/KO^*mice, only 2.1% of caput epididymal sperm were high quality (rating 7), in comparison to 22% of cauda sperm (p = 0.028, Fig. 4D-E). A comparison between genotypes and epididymis regions revealed that CCDC112 function is required to produce normal numbers of high-quality cauda epididymal sperm, with 63.3% being present in the *Ccdc112^WT/WT^* cauda epididymis versus 22% in *Ccdc112^KO/KO^* (p < 0.0001; Fig. 4D-E). Only 20.9% of caput sperm were classified as normal (rating 7) in *Ccdc112^WT/WT^*and 2.1% in *Ccdc112^KO/KO^* (p = 0.026; Fig. 4D-E).

The results indicate that, CCDC112 is required for the production of high-quality sperm midpiece structure. Notably, the magnitude and severity of the defects measured in sperm from *Ccdc112^KO/KO^* mice was significantly greater than that seen in wildtype males in both the caput and cauda epididymis. For *Ccdc112^KO/KO^* males, caput epididymal sperm, a significantly lower midpiece average normality rating (ranging from 1 – highly abnormal to 7 – high quality [normal]) of 3.75 was observed, compared to 5.07 in wildtype (p = 0.0033, Fig. 4G). For cauda epididymal sperm, an average normality rating of 5.38 was observed in *Ccdc112^KO/KO^*mouse sperm compared to 6.46 in wildtype (p = 0.036, Fig. 4G). Further, regardless of genotype, cauda epididymal sperm had more normal midpieces than those found in the caput, and accordingly, a significant increase in normality rating (*Ccdc112^WT/WT^* – 5.07 in caput to 6.46 in cauda, p = 0.0029; *Ccdc112^KO/KO^*– 3.75 in caput to 5.38 to cauda, p = 0.0008, Fig. 4G).

These data raise two possibilities: that the structure of the sperm midpiece continues to mature during epididymal transit, and/or abnormal sperm are removed from the epididymis during transit. To explore these possibilities, we measured sperm numbers in the caput and cauda epididymis in wildtype and *Ccdc112^KO/KO^* mice. As above, a 13% decrease in DSP was detected in the testis of *Ccdc112^KO/KO^* mice compared to wildtype littermates (p < 0.05, Fig. 1D). Significantly, in *Ccdc112^KO/KO^* individuals, a 54% decrease in sperm number was measured in the caput epididymis relative to wildtype (p < 0.0001, Fig. 4H), and a 35% decrease in sperm number in the cauda epididymis (p = 0.0053, Fig. 4H).

Collectively, these data reveal that as sperm transit through the epididymis (1) the midpiece continues to mature and (2) sperm with abnormal morphologies are selectively removed and act as a means of quality control. As an illustration of the later, sperm with large gaps absent of mitochondria and sperm completely devoid of mitochondria were observed in the caput of *Ccdc112^KO/KO^* individuals. As it is impossible for sperm to acquire additional mitochondria during epididymal transit, the absence of mitochondria-less sperm in the cauda epididymis is strongly indicative of their removal from the population. Equally, the presence of only 2.1% normal (rating 7) sperm in the caput epididymis *Ccdc112^KO/KO^* individuals compared to 22% normal sperm in cauda epididymis is conclusive of the ongoing maturation of the sperm midpiece during epididymal transit. Despite this quality control mechanism, the majority of cauda epididymal sperm from *Ccdc112^KO/KO^* males were abnormal, i.e. 22% of null sperm were morphological normal and only 15.6% were capable of progressive motility, compared to 63.3% and 50.2%, respectively, in sperm from wildtype. No sperm from *Ccdc112^KO/KO^* males could achieve fertility *in vivo*. Equally relative sperm numbers in the epididymis between genotypes indicate that abnormal sperm removal can occur in both the caput and cauda regions, and in the absence of CCDC112 culminate in a 35% reduction in overall sperm number. The existence of morphologically abnormal sperm in the caput epididymis in the absence of CCDC112, however, suggests that this quality control mechanism is not absolute and or it can be overwhelmed.

### CCDC112 is critical for normal flagella kinematics

To define how CCDC112 loss affected sperm tail waveform, high speed imaging analysis was performed on cauda epididymal sperm (53). As shown in Supplementary Video 1 and Figure 5, wildtype mouse sperm flagellum movement was flexible throughout all tail segments (Fig. 5A) and highly repetitive (Fig. 5B). By contrast, *Ccdc112^KO/KO^*mouse sperm flagellum movement was more rigid, notably in the midpiece (Supplementary Video 2; Fig. 5). An analysis of the flagella waveform revealed that the majority of sperm from *Ccdc112^KO/KO^* males possessed a stiff midpiece (Fig. 5A). Specifically, 17% of sperm from *Ccdc112^WT/WT^* mice had a highly flexible waveform wherein the amplitude of midpiece movement ranged between 27 – 39 µm (Fig. 5A), 60% were relatively flexible with a midpiece amplitude ranging 16 – 27 µm (Fig. 5A), 19% were moderately stiff 6 – 15 µm (Fig. 5A), and 4% were highly stiff with a midpiece amplitude ranging between 0.6 – 6 µm (data not shown). By contrast for *Ccdc112^KO/KO^* males, there were no highly flexible sperm, 12% of midpieces were relatively flexible (Fig. 5A), 28% were moderately stiff (Fig. 5A) and the majority of midpieces, 60%, were highly stiff (Fig. 5A). Collectively, 88% of sperm midpieces from *Ccdc112^KO/KO^* males were classified as stiff compared to 22% in wildtype. Correspondingly, the amplitude in the principal piece was significantly decreased (34.04 µm versus 16,79 µm; p = 0.0086, Fig. 5E) in *Ccdc112^KO/KO^*mouse sperm compared to their wildtype counterparts. The tail beating frequency (*i.e.*, the number of beating cycles completed per second) was comparable between genotypes (Fig. 5D).

**Figure 5.**
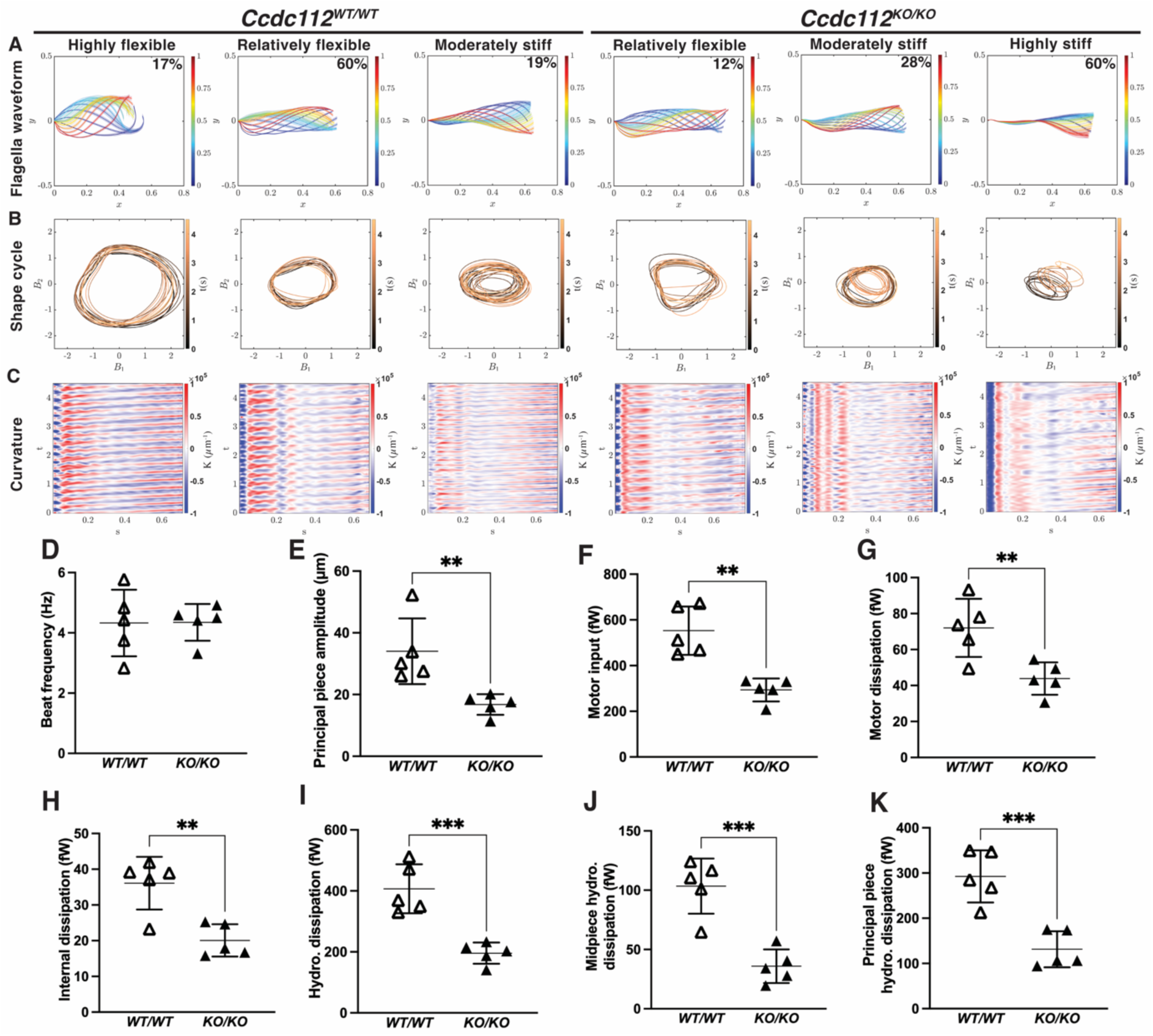
Sperm tail beating patterns and energetics are critically dependent on CCDC112. (A) Representative sperm flagella waveform in sperm from *Ccdc112^WT/WT^* and *Ccdc112^KO/KO^* mice over the proximal most 60 µm of the sperm tail. Coloured curves represent the non-dimensional time scale, where the beginning of the beat cycle is blue in the waveform and the end is in red. ‘y’ denotes the normalised amplitude and ‘x’ denotes the normalised length of the sperm tail where the y-intercept of 0 represents the sperm head. Percentage of sperm per genotype displaying each beat cycle type is denoted in the top right corner of each graph. (B) Representative shape cycles of the same sperm population in (A), as visualised by plotting the two dominant shape modes coefficients (B_1_ and B_2_) against each other over time (n = 5 animals/genotype and 8-15 sperm/animal). (C) Representative flagellum curvature plots, where the red and blue waves denote the direction of the flagellar bend above or below the sperm head hook and and the intensity of the colour denotes smaller radius of curvature (n = 5 animals/genotype and 8-15 sperm/animal). ‘T’ denotes time (seconds), and ‘s’ denotes arclength (µm/100) where 0 represents the sperm head. Beat frequency (D), principal piece amplitude (E), motor input (F), motor dissipation (G), internal dissipation (H), hydrodynamic dissipation (I), midpiece hydrodynamic dissipation (J) and principal piece hydrodynamic dissipation (K). Lines denote mean ± SD in D-K; ** P < 0.01, *** P < 0.001.

Next, to analyse waveform periodicity over time, sperm flagella shape cycle was assessed. Here, a circular shape cycle is indicative of highly reproducible and periodic waveform beating behaviour with a shape cycle displaying a more imperfect and disturbed circle indicating a more irregular beating behaviour. In wildtype mouse sperm, majority of the shape cycles were uniform circles with some irregularity being observed in sperm with more stiffened midpieces. Comparatively, *Ccdc112^KO/KO^* mouse sperm shape cycles presented most commonly with highly irregular, non-uniform circles indicative of significant variability in waveform between sequential beats.

To illustrate the role of CCDC112 in flagella, particularly midpiece flexibility over multiple beat cycles, we plotted the sperm tail curvature along the length of the tail over a period of 4.5 seconds (Fig. 5C). As shown for wildtype sperm, the alternating blue and red strips in the curvature kymograph indicate the wave propagation and formation along the flagellum is highly periodic and reproducible. By comparison, sperm from *Ccdc112^KO/KO^* males, particularly those with a highly stiffened midpiece, displayed considerably variable and disturbed flagella wave propagation and formation over both space and time (Fig. 5C). Collectively, these data demonstrate that CCDC112 plays a role in establishing sperm midpiece flexibility. The negative effect of CCDC112 loss on tail flexibility, and thus sperm tail amplitude was, however, observed in all regions of the tail.

At a subcellular level, sperm flagellum movement is generated by forces produced from dynein motors within the axoneme. Specifically, the power exerted from the flagellum dynein motors to the microtubules of the axoneme is termed the motor input (54). Our results showed that motor input was significantly lower in *Ccdc112^KO/KO^*mouse sperm in comparison to wildtype (561 fW versus 293 fW; p = 0.0011, Fig. 5F), indicating lower rate of energy production. Next, internal, and external dissipated power were assessed. Internal energy dissipation in the flagellum involves both motor dissipation, originating from the work done by other flagellar structures on dynein motors, and internal dissipation, arising from friction between cross-sectional planes within the flagellum. External energy dissipation, known as hydrodynamic dissipation, results from the friction between the flagellum and the surrounding fluid. Our data showed that all power dissipations were significantly reduced in *Ccdc112^KO/KO^*mice (p = 0.0093, Fig. 5G; p = 0.0033, Fig. 5H and p = 0.0007, Fig. 5I). Moreover, total hydrodynamic dissipation can be further divided into contributions from the midpiece and principal piece, both of which were significantly lower in sperm from *Ccdc112^KO/KO^* males (p = 0.005 and p = 0.0009, respectively, Fig. 5J-K).

### CCDC112 is required for optimal sperm metabolism

Ultimately sperm motility is driven by ATP hydrolysis by the dynein arms of the axoneme leading to microtubule sliding (55, 56). Within sperm, ATP production occurs via two dominant processes, oxidative phosphorylation in the mitochondria within the midpiece, and glycolysis via enzymes anchored to the fibrous sheath in the principal piece (57). To test the effect of CCDC112 loss and poor mitochondrial packing on mitochondrial respiration capacity (oxidative phosphorylation), we used the Seahorse mitochondrial stress test assay on both non-capacitated and capacitated sperm in wildtype versus *Ccdc112^KO/KO^*mice (Fig. 6). In support of previous reports demonstrating glycolysis to be the primary method of energy production in mouse sperm (6, 22, 58–60), our data showed no significant difference between mitochondrial respiration rates in non-capacitated versus capacitated wildtype mouse sperm (Fig. S5A). Similarly, in *Ccdc112^KO/KO^* males no significant difference between mitochondrial respiration rates in non-capacitated versus capacitated sperm was observed (Fig. S5B). When compared between genotypes, oxygen consumption data revealed that, in non-capacitated sperm, basal and maximal mitochondria respiration of sperm from *Ccdc112^KO/KO^* males was significantly impaired relative to their wildtype counterparts (34% basal and 45% maximal rate reduction; p < 0.01, Fig. 6A). Further, in capacitated sperm, both basal and maximal mitochondria respiration rates of sperm were also significantly diminished in *Ccdc112^KO/KO^* males compared to wildtype (45% basal and 50% maximal rate reduction; p < 0.01, Fig. 6B). CCDC112 function is thus required for optimal oxidative phosphorylation in sperm. We hypothesis this is via it’s essential role in mitochondrial sheath formation.

**Figure 6.**
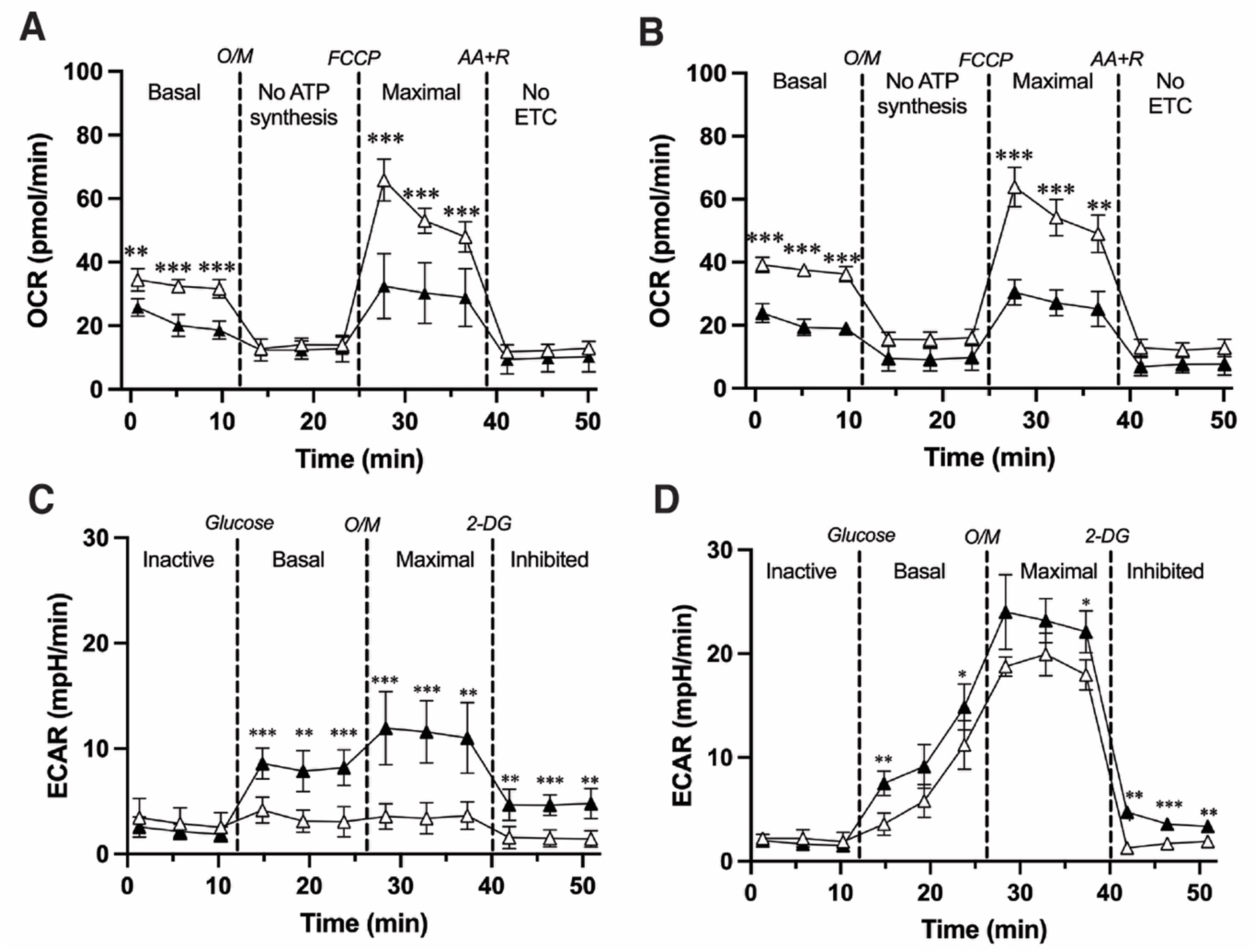
CCDC112 is essential for normal sperm energy production. Mitochondria stress test assay on non-capacitated (A) and capacitated (B) *Ccdc112^WT/WT^* (white triangles) and *Ccdc112^KO/KO^* mouse sperm (black triangles) (n = 3 mice/genotype). Glycolytic flux Seahorse assay on non-capacitated (C) and capacitated (D) *Ccdc112^WT/WT^* and *Ccdc112^KO/KO^*mouse sperm (n = 3 mice/genotype). Note the difference in the Y axis scale between mitochondrial stress test and glycolytic assays. OCR = oxygen consumption rate; ECAR = extracellular acidification rate; O/M = oligomycin; AA+R = antimycin A and rotenone; 2-DG = 2-Deoxy-D-Glucose. Lines denote mean ± SD; * P < 0.05, ** P < 0.01, *** P < 0.001.

As has been reported previously, capacitation in mouse sperm is associated with a notable increase in glycolytic activity as measured using a glycolytic flux Seahorse assay (24). In support, we observed a significant increase in the basal and maximal glycolysis activity in capacitated versus non-capacitated sperm for both wildtype and *Ccdc112^KO/KO^* males (p < 0.001 and p < 0.0001, respectively, Fig. S4C-D). When compared between genotypes and by contrast to the data for oxidative phosphorylation, sperm from *Ccdc112^KO/KO^*males exhibited significantly increased glycolysis activity (Fig. 6C-D). In the absence of CCDC112, the basal and maximal glycolysis capacity in non-capacitated sperm increased by 138% and 226%, respectively, compared to wildtype (p < 0.01, Fig. 6C) suggesting a degree of metabolic reprogramming in sperm lacking CCDC112. While in capacitated *Ccdc112^KO/KO^* mouse sperm, basal and maximal glycolytic activity increased by 53% and 22% relative to wildtype, (p < 0.05, Fig. 6D). As expected, when a competitive inhibitor, 2-deoxyglucose was added, sperm from wildtype mice returned to inactive levels, however, for *Ccdc112^KO/KO^*mouse sperm maintained a significantly higher glycolytic capacity, independent of capacitation status.

Finally, we assessed whether abnormalities in mitochondrial sheath structure affected sperm mitochondrial membrane potential using the JC1 probe. When mitochondrial membrane potential is high, JC1 accumulates in mitochondria and emits a red fluorescence. By contrast, JC1 emits a green fluorescence when potential is low. Mitochondrial membrane potential was unchanged between *Ccdc112^WT/WT^* and *Ccdc112^KO/KO^* mouse sperm (p = 0.13, data not shown), suggesting sperm mitochondria were functional at a basic level.

Collectively, these data reveal that CCDC112 is required for normal ATP production levels in non-capacitated and capacitated sperm. Membrane potential suggests that the decrease is due to mitochondrial sheath structural defects. We cannot, however, preclude the possibility that the abnormal packing of mitochondria disrupts mitochondria-mitochondria linking and synergistic effects on mitochondria function. While mitochondrial fusion has not been observed, or explored, in the mitochondrial sheath, intermitochondrial linker structures between adjacent mitochondria have been identified (61). The role of these linkers remain unknown, however, their association with mitochondrial cristae suggest the potential to facilitate electrochemical coupling, energy flow and/or communication between mitochondria (61, 62); processes which could all be disrupted in the absence of CCDC112.

### CCDC112 is a component of the distal appendages of the mother centriole in somatic cells

Given previous proteomics data suggested CCDC112 is a centriolar satellite protein in RPE-1 cells (47), we aimed to resolve CCDC112 localisation in somatic ciliated IMCD-3 cells by expressing exogenous eGFP tagged CCDC112. CCDC112 localisation was observed at the centrosome as marked by γ-tubulin (Fig. 7A). To refine localisation, cells were co-stained ODF2, a marker of the distal and subdistal appendages at the proximal end of the mother centriole (63). STED microscopy revealed that CCDC112 co-localised with ODF2 in the mother centriole (Fig. 7B). These findings were validated by co-staining with CEP164, a specific marker of distal appendages (64) and CEP170, a specific marker of subdistal appendages (65) revealing the localisation of CCDC112 solely to the distal appendages, and not subdistal appendages, in interphase cells (Fig. 7C).

**Figure 7.**
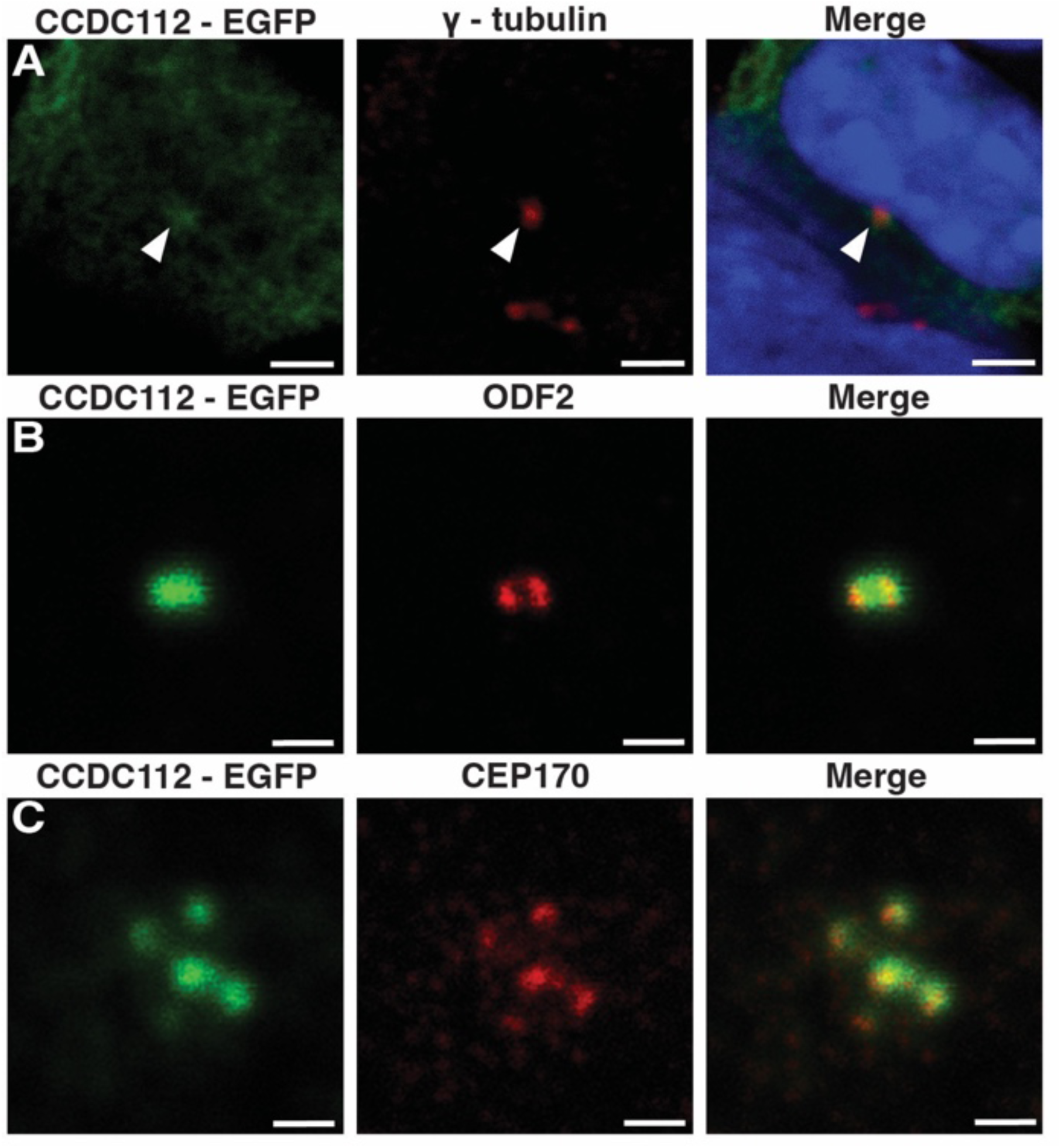
CCDC112 is a component of the distal appendages of the mother centriole. Immunofluorescence staining of somatic IMCD3 cells induced to produce a primary cilia via serum starvation expressing E-GFP tagged CCDC112 (green). CCDC112 localised to the centrosome (as marked by γ-tubulin; red) (A). Staining of the distal and subdistal appendages as marked by ODF2 (red) refined localisation to the mother centriole (B). Further staining localised CCDC112 specifically to the distal appendages (as marked by CEP170; red) of the mother centriole in interphase cells (C). Blue represents nuclei labelled by DAPI. Scale bars in A = 3 µm and in B-C = 1 µm.

Collectively, such a localisation and the reduced sperm tail length in a sub-population of sperm in mature *Ccdc112^KO/KO^* mouse sperm, indicate that CCDC112 is required for both normal motile cilia/flagella production by haploid germ cells, and for the assembly of the mitochondrial sheath.

## Discussion

The mitochondrial sheath is the defining structure of the sperm midpiece and abnormalities have been associated with human male infertility (31, 66–68). It provides structural support and hypothesized to generate energy necessary for multiple aspects of flagella movement (3, 4, 6). While many of the key steps of sperm midpiece formation have been described at a cytological level (31), the molecular processes required for assembly and function remain poorly understood. Herein, we show that CCDC112 is an essential for both processes. Specifically, we have identified a previously unrecognised process of mitochondrial sheath maturation that occurs during epididymal sperm maturation. We demonstrate a critical role for CCDC112 in mitochondrial morphogenesis and remodelling during mitochondrial sheath formation in the testis and remodelling in the epididymis. We show that such remodelling and the associated optimisation of ATP production is critical to the manifestation of a highly effective sperm flagella waveform and mechanical power and, in turn, the capacity of sperm to reach and fertilise oocytes within the female reproductive tract.

Our data establish the novel process of mitochondrial sheath maturation as one of the key events in the sperm epididymal maturation process. While other types of sperm maturation within the epididymis have been well documented, including biochemical and plasma membrane modifications (69–72), the acquisition of potential for sperm motility (73) and the transfer of RNA cargoes, proteins and lipids via sperm-epididymosomes (epididymis specific extracellular vesicles) interactions (74), no study to date has described complex sperm structural modifications during this period. Our data show that at the time of spermiation, the majority of spermatozoa are structurally immature, and that maturation continues as sperm transit through the epididymis from the caput to the cauda. Such maturation includes elongation/morphogenesis of mitochondria into a rod-like form and/or further compaction of mitochondria within the sheath. In agreement, we observed an increase in sperm population midpiece quality in the distal epididymis. Simultaneously, our data evidence that poor-quality sperm with highly abnormal midpiece phenotypes e.g. absent mitochondria, are selectively removed from the epididymis during transit.

While the specific removal of sperm with internal structural defects is a novel observation, the selective removal of abnormal sperm is consistent with a report from Glover (75), who observed a higher rate of decapitated and dead sperm in the proximal corpus compared to the cauda in cat, dog, goat and bovine epididymides. While the exact method of removal remains poorly defined, data from numerous mammalian species, including mice, suggest that misshapen sperm are captured in a dense matrix formed via the release and subsequent merging of aposomes released from the epididymal epithelium (76–78). The entanglement of abnormal sperm separates them from viable sperm, allowing for their disintegration and dissolution by epididymosomes, present within aposomes (76). Further, while instances of sperm phagocytosis or spermiophagy have been reported in mammals (79–81), this is a rare occurrence and is not the primary quality control mechanism mediating the removal of abnormal sperm (82, 83). Data presented here also suggest that while this quality control process can scale to some degree, as indicated by the 35% reduction in *Ccdc112^KO/KO^* cauda epididymal sperm count compared to wildtype, at some point it is overwhelmed, leading to the presence of residual abnormal sperm in the ejaculate. This selective removal of abnormal sperm sits ‘on top of’ an analogous process that occurs in the testis that can trigger spermiation failure (84, 85).

As defined above, mitochondrial loading and remodelling within the midpiece is a multi-step process. In the absence of CCDC112, the recruitment of mitochondria from the cytoplasm appears largely normal (Fig. 8). The translocation and attachment of mitochondria to the outer dense fibres also appears mostly unaffected, with the exception of a small percentage of cells where that mitochondria have failed to load onto, or attach to, the outer dense fibres. By contrast, the morphogenesis of mitochondria from spherical to crescent then to rod shaped and the staggering and compaction of mitochondria into the distinct double helical structure of the sheath, was impaired in the absence of CCDC112. A result of these defects, and an underlying deficit in ATP production, sperm presented with a stiffened midpiece with limited flexibility. Consequently, in the absence of CCDC112 the flagellar waveform was unable to produce sufficient power to sustain progressive motility to reach the upper oviduct and fertilise oocytes. In summary, our data suggest a role for CCDC112 in regulating mitochondrial sheath formation and potentially in the optimisation of ATP production via oxidative phosphorylation and maintaining midpiece flexibility.

**Figure 8.**
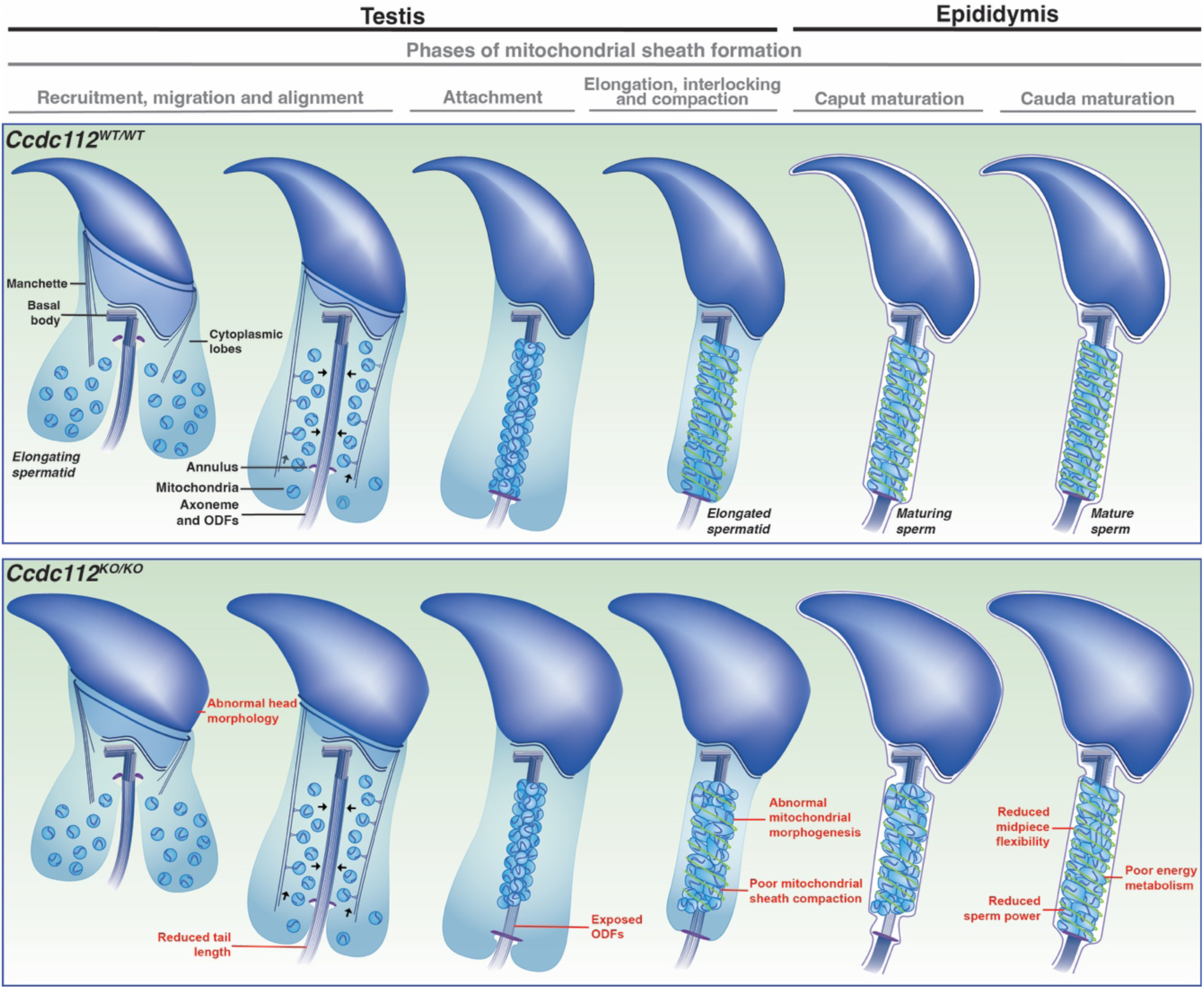
Summary schematic of sperm formation in the absence of CCDC112. In *Ccdc112^KO/KO^* mice, sperm commonly possessed abnormal mitochondrial sheath structure with overall irregular mitochondrial morphology and poor mitochondrial elongation, staggering and compaction during sheath formation. A subset of sperm also possessed exposed outer dense fibres (ODFs) and immature mitochondria within the sheath, abnormal head morphology and shortened tails. As a result, sperm exhibited poor energy generation, reduced tail flexibility, particularly in the midpiece, and reduced sperm power. Image modified from (55).

Our data also support a facilitative role for CCDC112 as a component of the distal appendages of the basal body, in the development of the sperm axoneme and thus sperm tail length. In mouse spermatogenesis the basal body docks to both the sperm nucleus and plasma membrane in step 2/3 (86). This ultimately forms a separate lobe, the ciliary lobe, which is isolated from the remainder of the cytoplasm (the cytoplasmic lobe) (2). Subsequently, the axoneme extends from the basal body into the ciliary lobe to form the core of the sperm tail (2). The localisation of CCDC112 to the distal appendages in IMCD-3 cells suggests it functions at a similar location in round spermatids, where it may ultimately contribute to controlling protein entry into the ciliary compartment. The hypothesis is supported by the presence of a distinct population of short sperm within the cauda epididymis from *Ccdc112^KO/KO^* males but remains to be formally tested. Whether such abnormalities in either sperm tail length and/or mitochondrial sheath structure are the cause of abnormal retention of elongated spermatozoa in the testis also remains to be tested.

In summary, we propose that CCDC112 executes its function via two biological processes. First, CCDC112 may function at the basal body to facilitate early steps of tail development by contributing to the efficient regulation of protein entry into the sperm tail (87). Second, CCDC112 directly contributes to tail function through a role in mitochondrial sheath formation and in optimising ATP production. Collectively, this study adds to the field’s knowledge of how functional sperm are produced and the causes of male infertility. We reveal a novel form of epididymal sperm maturation, a novel mechanism of mitochondria-mitochondria interactions of potential relevance to multiple tissues, and the complex relationship between energy production and sperm tail kinematics.

## Materials and Methods

### Knockout mouse model production

All animal procedures were conducted in accordance with the animal ethics guidelines generated by the Australian National Health and Medical Research Council (NHMRC). Experimental procedures were approved by The University of Melbourne Animal Ethics Committee (application 20640) or Monash University Animal Experimentation Ethics Committee (BSCI/2017/31).

A *Ccdc112* knockout mouse line was generated on the C57BL/6J background through the Monash University Genome Modification Platform (a partner of the Australian Phenomics Network) using CRISPR/Cas9 technology. Guide RNAs targeting regions upstream and downstream of exon 2 (GCAGTGACCGCGCATGCACATGG and TTCCTAATACTGTGAGACTGGGG, respectively) and Cas9 machinery were injected into fertilised mouse eggs (zygotes) to target the removal of exon 2 of *Ccdc112* of the principal (longest) transcript (*ENSMUST00000072835.7* [2801 bp]). The 122 bp deletion was confirmed via Sanger sequencing. This modification led to a premature stop codon in exon 3 of *Ccdc112* and a non-functional product. Mice heterozygous for the *Ccdc112* deletion were bred to generate knockout individuals (*Ccdc112^KO/KO^*) and wildtype controls (*Ccdc112^WT/WT^*). Genotyping was performed using the primers (Forward – 5’-GTTCGGAACAGTGTGCGGAG-‘3 and Reverse – 5’-AGCTGGCTTGAGTTCTGCTT-‘3) which amplified a 1050 bp wildtype band and 680 bp knockout band. In the effect of exon deletion on Ccdc112 mRNA expression was assessed using quantitative polymerase chain reaction (qPCR) on testis cDNA primers spanning exons 2 to 4 of the two longest transcripts (*ENSMUST00000072835.7* [2801 bp] and ENSMUST00000238189.2 [761 bp]) (Forward – 5’-GCTGTAGTGCTTCAGGTGG-‘3 and Reverse – 5’-AGTCCCTCTTTGGTTGTAG-‘3). qPCRs were performed using SYBR Select Master Mix (Applied Biosystems). Each reaction was performed in triplicate for wildtype and quadruplicate for *Ccdc112^KO/KO^* using the Agilent Mx3000P qPCR system or an Applied Biosystems QuantStudio 3 Real-Time PCR System with the parameters of 50°C for 2 minutes, 95°C for 10 minutes followed by 40 cycles of 95°C for 15 seconds and 60°C for 1 minute. *Ppia* was amplified at the same time as a housekeeping control (using 5’-CGTCTCCTTCGAGCTGTTT-’3 forward and 5’-CCCTGGCACATGAATCCT-’3 reverse primers), and all results were normalised to *Ppia* expression. Differential expression was analysed using the 2ΔΔCT method (88).

### Analysis of *Ccdc112* expression

To investigate the expression of *Ccdc112* in human tissue, we obtained single cell RNA sequencing data from (50). To assess expression in human and wildtype mouse testis cells we obtained single cell RNA sequencing data from (51) and (52), respectively. Usage of these data sets was done in accordance to the Creative Commons Attribution 4.0 International License (http://creativecommons.org/licenses/by/4.0/).

### Fertility characterisation

Male fertility in *Ccdc112* knockout male mice was assessed as per the framework described in (89). Briefly, knockout male mice at 10-12 weeks of age were mated with two wildtype females (40-80 days of age). Successful mating was confirmed by the presence of copulatory plugs, and litter size was recorded as the number of pups born per plug. Males were subsequently culled and weighed. For histological examination, testes and epididymides were dissected, weighed, and fixed in Bouin’s fixative for 5 hours, processed into paraffin blocks, and cut into 5 µm sections using standard methods. For sperm motility assessment, cauda epididymal sperm were backflushed into MT6 medium at 37 °C, allowed to swim out for 15 min, then examined via computer assisted semen analysis (‘CASA’) (n ≥ 1000 cells/animal; n ≥ 3 animals/genotype) (90, 91). To assess if a membrane permeable energy source could rescue sperm motility, MT6 medium was supplemented with 55 µg/ml adenosine 5’-triphosphate (ATP) disodium salt hydrate (A2383, Sigma, USA) and CASA analysis was repeated (92). Sperm from the base MT6 solution (without ATP supplementation) were subsequently washed in PBS and airdried onto SuperFrost slides for morphological analysis.

Periodic acid-Schiff’s reagent and haematoxylin staining was undertaken and used to visualise testis histology (n ≥ 5/genotype), while epididymis histology and sperm morphology were investigated using haematoxylin and eosin staining (n ≥ 3/genotype). Epididymal sperm morphology was assessed and broadly categorised as “normal” as determined by the most common sperm phenotype observed in wildtype mice or “abnormal”, then further categorised into cellular defects, including “abnormal sperm head morphology”, “abnormal sperm midpiece structure”, “short tail” and/or “coiled tail” (n ≥ 100 sperm/animal; n = 3 animals/genotype). Midpiece and tail length were also measured using Fiji (version 2.9.0/1.53t) (n = 5 animals/genotype; n ≥ 100 sperm/animal for tail length and n ≥ 20 sperm/animal for midpiece length).

Testis daily sperm production and epididymal sperm content were determined on snap frozen tissues using the Triton X-100 nuclear solubilisation method as previously described in (93) on mice aged 70-84 days (n ≥ 3/genotype). Quantification of round spermatid number per stage VIII tubule was conducted across 10 tubule cross-sections wherein the cross-section was round as an indication of dissection perpendicular to the length of the tubule. Tubule area was measured using Fiji (version 2.9.0/1.53t) (n = 5 animals/genotype). Germ cell apoptosis was examined via immunostaining for cleaved caspase 3 and 7 as outlined in (93). The number of caspase positive cells was counted in a minimum of 100 randomly selected tubules per mouse (n = 5 animals/genotype).

To assess the ability of sperm to traverse the female reproductive tract to reach the oviduct, wildtype and *Ccdc112^KO/KO^*male mice were plug mated overnight with wildtype females. Before 9 am plugs were checked and upon detection of a positive plug females were culled at 10 am (an estimated 10 hours post-mating). Oviducts were dissected to separate regions of the isthmuses and ampullae. Each pair of oviduct regions was then minced and dried onto a glass slide after removing large chunks of tissue, then fixed with 4% paraformaldehyde in PBS (10 mM Na_3_PO_4_, 2.68 mM KCl, 140 mM NaCl) for 10 minutes before being washed in PBS. Sections were stained with haematoxylin and eosin and the number of sperm per region counted (n ≥ 3/genotype).

### Electron microscopy

To investigate caput and cauda sperm mitochondrial sheath structure via scanning electron microscopy, sperm were isolated from the caput epididymis by careful dissection and nicking of the cauda epididymis with a sharp scalpel blade in warm PBS in a Petri dish at 37 °C for 30 min. For the cauda epididymis, sperm were collected using the backflushing method (94), then incubated in 100 µl of 1 × PBS (10 mM Na_3_PO_4_, 2.68 mM KCl, 140 mM NaCl) for 30 min to remove of the plasma membrane (95). Scanning electron microscopy was undertaken as described in (95). To investigate cauda epididymis sperm ultrastructure via transmission election microscopy, cauda epididymal sperm were backflushed into MT6 medium before being processed as previously described (93). Images were captured on a Talos L120C for TEM or a FEI Teneo VolumeScope for SEM at the Ian Holmes Imaging Centre (The University of Melbourne, Australia) or a Jeol 1400 Plus electron microscope for TEM at the Vera and Clive Ramaciotti Centre for Electron Microscopy (Monash University, Australia).

### Scoring of mitochondrial sheath normality

After noticing abnormalities in mitochondrial sheath structure in sperm from *Ccdc112^KO/KO^* males, the degree of sheath abnormality was quantified on SEM images (n ≥ 5 mice/genotype). For each biological replicate, a minimum of 10 sperm from each region of the epididymis were assessed and scored from 1 [highly abnormal] to 7 [normal]. We defined these categories as: 1 – mitochondria were completely absent or the mitochondrial sheath was stratified (highly elongated mitochondria morphed together); 2 – mitochondria were frequently immature, morphed together, incorrectly oriented or sections of the sheath lacked mitochondria; 3 – mitochondria were often immature (round/crescent), morphed together and/or incorrectly oriented; 4 – many mitochondria had abnormal orientations and size, including mitochondria morphed together (beginning and end of mitochondria was not discernible); 5 – mitochondria were loosely coiled with some incorrectly oriented (more horizontal, or diagonal orientation – i.e., less perpendicular to the axoneme) and heterogeneously sized mitochondria; 6 – mitochondria were less tightly coiled and/or a small section of the sheath bearing mitochondria had misaligned ends; 7 – no defects, with mitochondria in the sheath being tightly packed, regularly coiled, homogenously sized and correctly orientated (i.e., almost perpendicular to the axoneme). Using a strict rating system, a score of 1-6 was considered abnormal i.e. displayed disordered mitochondrial packing. To ensure the rating system used was unbiased and reproducible, two biological replicates for each genotype were blindly assessed and rated by a second researcher and found to match the original score. To further define the disordered mitochondrial sheath packing of caput sperm, defects were separately classified to define the incidence of immature mitochondria (spherical and/or crescent shaped), exposed outer dense fibres and/or no mitochondria.

### High resolution sperm motility analysis

To assess the kinematics of sperm flagellar movement, beating patterns of head-tethered cauda epididymal sperm were recorded and analysed at high resolution. Video capture and analysis was undertaken as outlined previously (53). Briefly, sperm were backflushed from the cauda epididymis into modified TYH medium (10 mM HEPES, 5.6 mM glucose, 1 mM MgSO_4_, 1.2mM KH_2_PO_4_, 2 mM CaCl_2_, 4.8 mM KCl, 135 mM NaCl, 0.5 mM sodium pyruvate, 10 mM sodium L-lactate), supplemented with 0.3 mg/ml of bovine serum albumin at 37 °C. After 15 min, sperm were diluted 1 in 10 in TYH medium. A custom-made imaging chamber was made by affixing two strips of clear adhesive tape 15 mm apart on a glass microscope slide. 10 µl of the diluted sperm suspension was added in between the tape and covered with a glass coverslip (Menzel Gläser). Sperm motility patterns were then imaged using an Olympus AX-70 upright microscope outfitted with a U-DFA 18 mm internal diameter dark-field annulus and a 20× 0.7 NA objective (UPlanAPO, Olympus, Japan). Images were captured via a Zyla 5.5 CL10 sCMOS camera (Andor Technology, Northern Ireland) at 250 frames per second using the Fiji image-processing package with the Micro-Manger Studio plugin (version 1.4.23). Ten videos per animal (n = 5 animals/genotype) capturing sperm tail movement over 1000 frames were taken. Images were then adjusted and processed in Fiji (version 2.9.0/1.53t) before being analysed in MATLAB (version R2021b) using a modified version of the MATLAB codes used in (53) and publicly available from the Monash University Research Repository (DOI: 10.26180/14045816). Briefly, analysis of each image was conducted by extracting the sperm flagella centreline and reconstructing its waveform using an automated image analysis algorithm as previously described in (53). The curvature kymograph was plotted by calculating the tangent angle of the flagella waveform. The beating frequency of the sperm flagella was calculated by determining the average frequency of curvature turning points at 50% and 75% of the flagella length. The mean sperm flagella amplitude of the midpiece and principal piece was calculated by using set points corresponding to the sperm head (6.3 µm), mid (22.4 µm) and principal (80 µm) piece lengths (35) and determining the average amplitude along each tail section. Analysis of the flagella power (motor input power and dissipated powers) for each image, were calculated as previously described in (53). Briefly, motor input was defined as power expended from the dynein motors towards the sperm flagellum (54). Motor dissipation was defined as energy dissipated by the dynein motors due to work done of them from other flagella structures, internal dissipation was defined as energy dissipation due to friction between flagellum cross sectional planes and hydrodynamic dissipation was defined as energy dissipation into the fluid surrounding the flagellum.

Flagella waveform graphs were categorised into 4 main groups based on their midpiece amplitude. Midpieces with an amplitude ranging from 27 – 39 µm were considered highly flexible, 16 – 27 µm were relatively flexible, 6 – 15 µm were moderately stiff and 0.6 – 6 µm were highly stiff.

### *In vitro* fertilisation

To assess the ability of sperm to interact and fertilise oocytes, *in vitro* fertilisation was undertaken. To induce superovulation, 3-5 week old C57BL6/CBA (‘F1’) female mice were administered with 100 µl (5 IU) intraperitoneal injections of equine chronic gonadotrophin (Folligon) followed by 100 µl (5 IU) injections of human chorionic gonadotrophin (hCG – Chorulon) 48 hours later. The following day, male mice were culled, and their cauda epididymides dissected into warm mineral oil. Sperm were then backflushed gently into 200 µl pre-equilibrated drops of G-IVF PLUS (Vitrolife, Sweden) under OVOIL in 35 mm Petri dishes and incubated for 1 hour at 37 °C in 6% CO_2_ to capacitate. In parallel, 14 hours after the second round of injections, female mice were culled, and their ovaries were dissected and placed in 1 m G-IVF PLUS at 37 °C. Cumulus-oocyte complexes were retrieved from the ampulla and washed 3 times in high calcium HTF media (25 mM NaHCO_3_, 5.55 mM glucose, 0.2 mM MgSO_4_.7H_2_O, 0.31 mM KH_2_PO_4_, 5.14 mM CaCl_2_.2H_2_O, 4.69 mM KCl, 101.61 mM NaCl, 2.72 mM sodium pyruvate, 26 mM sodium L-lactate, pH 7.4) supplemented with 4 mg/ml bovine serum albumin (fraction V), 10 µl/ml penicillin-streptomycin and 1 mM reduced glutathione (GSH; G4251, Sigma). Complexes were then moved to a fertilisation dish of HTF media and GSH covered with OVOIL, to which 5 x 10^5^ sperm were added and incubated at 37 °C in 6% CO_2_, 5% O_2_ for 4 hours. Presumptive zygotes were washed in 50 µl drops of pre-equilibrated G1 PLUS medium in 35 mm Petri dishes at 37°C in 6% CO_2_ for 30 min, then cultured in G1 PLUS overnight. The following morning, the number of 2-cell embryos was counted, and 2-cell rate determined.

### Immunochemistry

To assess protein localisation in the paraffin embedded testis sections, sections were processed and stained as previously described using diaminobenzidine (DAB) as the chromogen (96).

To assess protein localisation in cultured cells, cells were fixed in cold 100% methanol, before non-specific antibody binding was blocked with CAS-block, incubated with primary and secondary antibodies, counterstained with DAPI and mounted, as above.

To examine oocytes 16 hours post-fertilisation, cells were fixed in 4% PFA for 30 mins, washed in PBS, permeabilised in 0.2% Trition-X for 1 hour. Following, cells were washed in PBS, stained with DAPI for 20 min, before being washed in PBS and mounted in µ-slide 8 well glass bottom plate (80827, Ibidi Gmbh, Germany).

Primary antibodies used included those against α-tubulin (T5168, Sigma; 0.1 µg/ml), CEP170 (ab84545, Abcam, USA; 1 in 500), cleaved-Caspase 3 (9664, Cell Signalling; 0.5 μg/ml), cleaved-Caspase 7 (9491, Cell Signalling, 0.23 μg/ml), GFP (11814460001, Roche, Switzerland; 0.4 µg/ml), ODF2 (HPA001874, Sigma; 0.28 µg/ml) and γ-tubulin (ab11317, Abcam, 11.7 µg/ml). Trialled commercial CCDC112 primary antibodies included HPA045120 (Sigma), HPA050621 (Sigma) and ab237742 (Abcam). Secondary antibodies were used diluted 1 in 500 and included Alexa Fluor 488 donkey anti-mouse (ab150105, Abcam) and Alexa Fluor 555 donkey anti-rabbit (ab150074, Abcam).

Brightfield images were collected on the Olympus BX53, Olympus DP80 camera with the Olympus Cell Sens Dimension software (v3.1.1). Immunofluorescent images were captured using either a Leica TCS SP8 confocal microscope (Leica Microsystems) or the Elyra LSMS880 (Zeiss, Germany) in the Biological Optical Microscopy Platform facility (University of Melbourne) or an Abberior STED-Super-Resolution Microscope (Abberior, Germany) at the Monash Micro Imaging Facility (Monash University). All immunofluorescent images were taken using the 63x/1.40 HC PL APO CS2 oil immersion objective besides oocyte images which were taken using the 20x/0.8 Plan-Apochromat air objective. Images were adjusted uniformly across the image and between groups.

### Mitochondrial membrane potential

Mitochondrial membrane potential was analysed using mitochondrial fluorescent probe JC-1 (T3168, Invitrogen) using a modified version of the analysis previous described in (97). Briefly, sperm were backflushed from the cauda epididymis into MT6 medium (excluding phenol red) at 37 °C. Sperm were then incubated with 1 µM JC1 in MT6 (T3168, Invitrogen) for 20 min at 37 °C in the dark. Simultaneously, LIVE/DEAD staining was performed (L7011, Invitrogen) by incubating cells with SYBR14 (1 µl/ml) and 5 µM propidium iodide (PI) for 10 min at 37 °C in the dark. To do so, PI and SYBR 14 dye was added halfway through JC-1 incubation. Following incubation, cells were centrifuged for 1 min at 300 × g and then gently resuspended in fresh MT6 medium before being imaged. To prevent sperm swimming out of frame during imaging, custom-made laminin coated observation chambers were created. To do this, two strips of clear tape were affixed at either end of a glass slide, before 0.2 mg/ml laminin (Thermo Fisher Scientific, USA) in TBS was added to slides and allowed to coat for 1 hour at 37 °C in a humified chamber. After incubation, slides were washed with water and kept warmed until used. Per slide, 100 – 150 µl of the sperm suspension was added in between the tape and covered with a glass coverslip (Menzel Gläser, thickness 1). Sperm were then imaged using filters green (FITC), red (CY3) and far-red (PI) on an Olympus BX53, Olympus DP80 camera with the Olympus Cell Sens Dimension software (v3.1.1). Mitochondrial membrane potential of sperm was then quantified on live cells (PI/far-red negative) by measuring the fluorescence intensity of the red channel (high membrane potential) relative to the green channel (low membrane potential) using Imaris (10.0.1), n ≥ 20 sperm/animal; n = 3 animals/genotype).

### Sperm mitochondrial respiration and glycolytic flux analysis

To assess sperm metabolism, Seahorse real-time metabolic analysis was undertaken. First, the assay cartridge was prepared. The day prior to the assay, the Seahorse sensor cartridge was hydrated in sterile water in a non-CO_2_ 37°C incubator overnight. Following hydration, the sensor cartridge was then calibrated via incubation in pre-warmed Seahorse XF Calibrant (incubated overnight in a non-CO_2_ 37°C incubator) for 45 – 60 minutes prior to loading the injection ports of the sensor cartridge. After incubation, the cartridge was removed from the incubator and each port was loaded as follows: For the mitochondrial stress test assay – 1^st^ injection port – 20mM Oligomycin A (75351, Sigma); 2^nd^ injection – 20uM FCCP (C2920, Sigma); 3^rd^ injection port – 10mM Rotenone (R8875, Sigma) and 10mM Antimycin A (A8674, Sigma) and 4^th^ injection – a modified TYH media ((2 mM HEPES, 5.6 mM glucose, 1.2 mM MgSO_4_, 1.2 mM KH_2_PO_4_, 1.7 mM CaCl_2_, 4.7 mM KCl, 138 mM NaCl), supplemented with 3 mg/ml of bovine serum albumin). For the glycolytic stress test assay – 1^st^ injection port – 100mM D-Glucose (A783, Univar, USA); 2^nd^ injection port – 20mM Oligomycin A; 3^rd^ injection port – 500mM 2-Deoxy-D-Glucose (L07338.06, Thermo Fisher Scientific) and 4^th^ injection port – a modified TYH media ((1 mM HEPES, 1.2 mM MgSO_4_, 1.2 mM KH_2_PO_4_, 1.7 mM CaCl_2_, 5.4 mM KCl, 144 mM NaCl), supplemented with 3 mg/ml of bovine serum albumin). These injections are then diluted 10 fold to leave the final concentrations for the mitochondrial stress test assay at 2mM oligomycin A, 2mM FCCP, 1mM Rotenone and 1mM Antimycin A and for the glycolytic stress test assay at 10mM D-Glucose, 2uM oligomycin A and 50mM 2-Deoxy-D-Glucose. The cartridge was then transferred to the XFe96/XF96 instrument and calibration via the Wave software (version 2.6.3) began. In parallel, to ensure sperm heads adhered to the Seahorse plate, each plate well was incubated with 30 µl of mouse laminin diluted with sterile filtered TBS (0.2 mg/ml; 23017015, Thermo Fischer Scientific) at 4°C overnight. The following day, the laminin coated Seahorse plate was incubated in a non-CO_2_ 37°C incubator for 1.5 hours. The majority of the laminin was then removed by pipetting, leaving a thin coat, and the plate rinsed with MilliQ water just prior to the addition of sperm. For the assay, cauda epididymal sperm were backflushed into 1 ml of a modified TYH medium at 37 °C. After 15 min, sperm in solution were transferred to a new pre-warmed microfuge tube. 1.2 million sperm per well were then added to the Seahorse plate. The plate was then spun with no braking in a prewarmed 37 °C centrifuge at 300 g for 5 min. Cells were than rested in a non-CO_2_ 37°C incubator for 15 minutes prior to starting the assay. The Seahorse plate was then loaded into the XFe96/XF96 instrument following Seahorse sensor cartridge calibration and analysis was conducted using the Wave software. At the end of the experiment the number of sperm per well was then re-counted using a haemocytometer to avoid bias from differential adhesion. Results were normalised accordingly.

### Cell culture and transfection

To examine the localisation of CCDC112 in cilia, IMCD3 cells (ATCC) were transfected with a CCDC112-EGFP vector. The expression vector was constructed by inserting a copy of the *Ccdc112* mouse cDNA (Forward – 5’-GTCTCGACGAGGAAACATGG-‘3 and Reverse – 5’-TAGGAACAGAACTGACAATGCTC-‘3), amplified from wildtype C57BL/6J testis cDNA by PCR reaction, into a pEGFP-C1 expression vector. IMCD3 cells were cultured in DMEM/F12 medium supplemented with 10% FBS and 1% penicillin and streptomycin. The CCDC112-EGFP vector was then transfected into IMCD3 cells using Lipofectamine 2000 Reagent (Thermo Fisher Scientific). Briefly, cells were seeded and incubated as per the manufacturer’s instructions. Ciliogenesis was then induced by culturing IMCD3 cells on coverslips in OptiMEM serum free medium (Thermo Fisher Scientific) for 48 hours as previously reported (98, 99). Cells were then fixed, stained, and imaged as detailed below.

### Statistics

Statistical analyses were conducted in GraphPad Prism 10 (version 10.0.3) with the statistical difference between genotypes determined using an unpaired student’s T-test on averages of technical replicate data per animal. The statistical difference between more than two groups was determined using a two-way ANOVA and Tukey post-hoc analysis, respectively. Significance was defined as *p* < 0.05. Specifically, statistical differences for breeding data, weights, DSP, ESC, round spermatids per tubule numbers, oviduct counts, CASA and high-resolution sperm motility analysis, high-resolution sperm power outputs, sperm midpiece and tail lengths, sperm morphology (brightfield), caput midpiece comparison, *in vitro* fertilisation, mitochondria Seahorse stress test assay, glycolytic flux Seahorse assay and mitochondrial membrane potential, an unpaired student’s T-test was used. For the remaining mitochondrial sheath abnormality comparisons (normal mitochondrial sheath data, rating of midpiece abnormality severity and relative sperm count) a two-way ANOVA was used. Statistical analysis of germ cell apoptosis data was conducted using generalised linear mixed models with zero-inflated negative binomial distribution as determined by the Akaike information criteria estimates (100). Analysis was undertaken in R version 4.1.10 (R Core Team 2021) and R Studio version 1.4.1717 (RStudio Team 2020) using the lme4 (101) and glmmTMB (102) packages.

## Acknowledgments

This research was funded by an Australian Research Council Discovery project (DP230100747), Bill and Melinda Gates Foundation Funding to MKOB and funding from the National Institutes of Health (R01HD078641 and P50HD096723). MLG is the recipient of a scholarship from the University of Melbourne (Australia) and RN acknowledges funding from the Australian National Health and Medical Research Council (NHMRC) Investigator Grant (2017370).

## Declaration of interests

The authors declare no competing interests.

**Figure S1.**
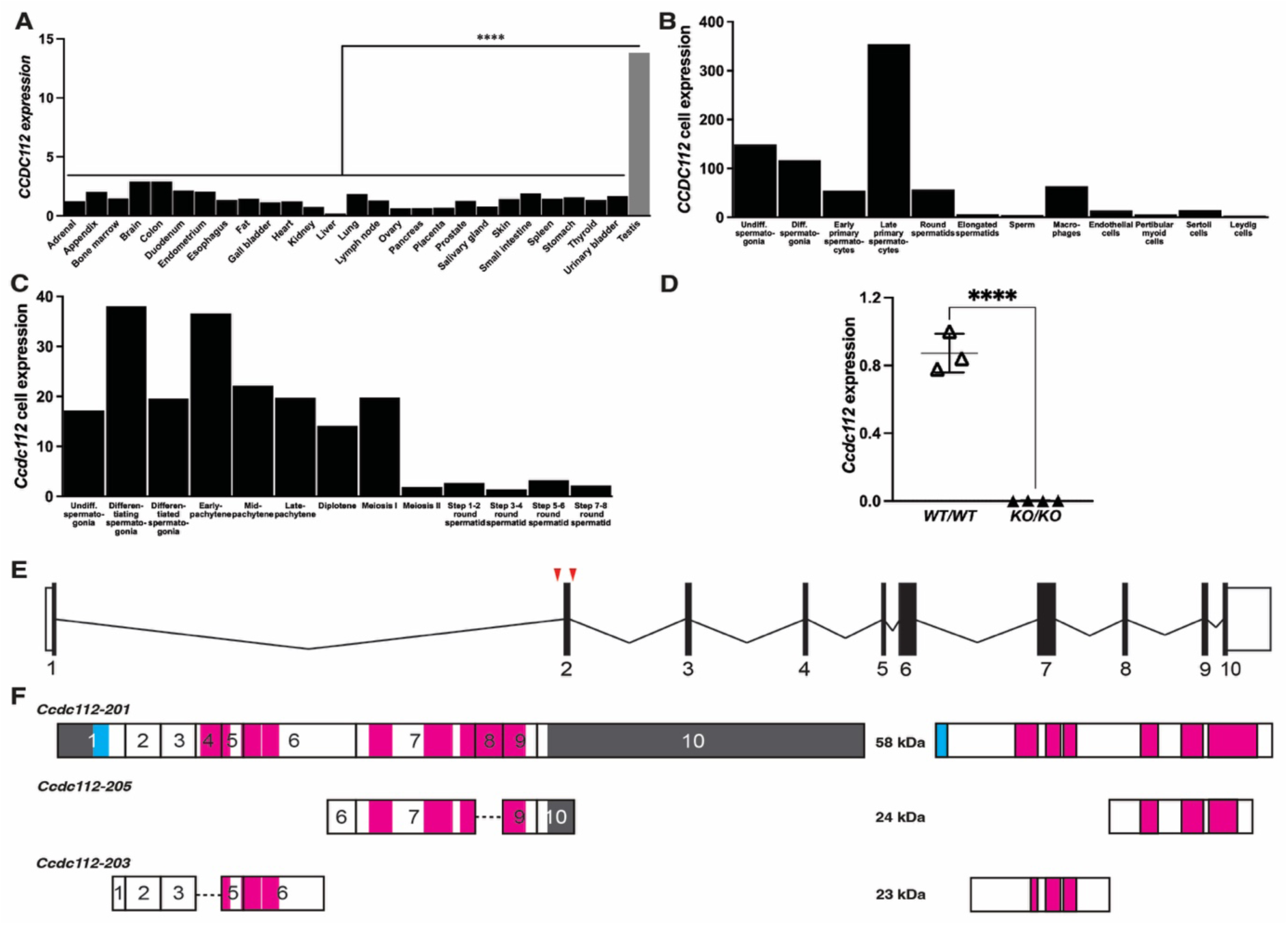
*Ccdc112* knockout mouse model and expression in male germ cells. *Ccdc112* expression in human organs (A) and human (B) and mouse (C) testis cell types as determined by single cell sequencing data (50–52). (D) *Ccdc112* expression in wildtype and knockout testes, as measured by qPCR (mean ± SD). Mouse *Ccdc112* exon map (E), including the predicted transcripts and their domains, and the corresponding proteins encoded (F). Signalling protein and coiled-coil domains are shown in blue and pink, respectively, while untranslated regions are shown in grey. Red arrowheads in (E) indicate guide RNA target regions used to excise exon 2 in the *Ccdc112^KO^*^/*KO*^ mouse line. Lines denote mean ± SD; **** P < 0.0001.

**Figure S2.**
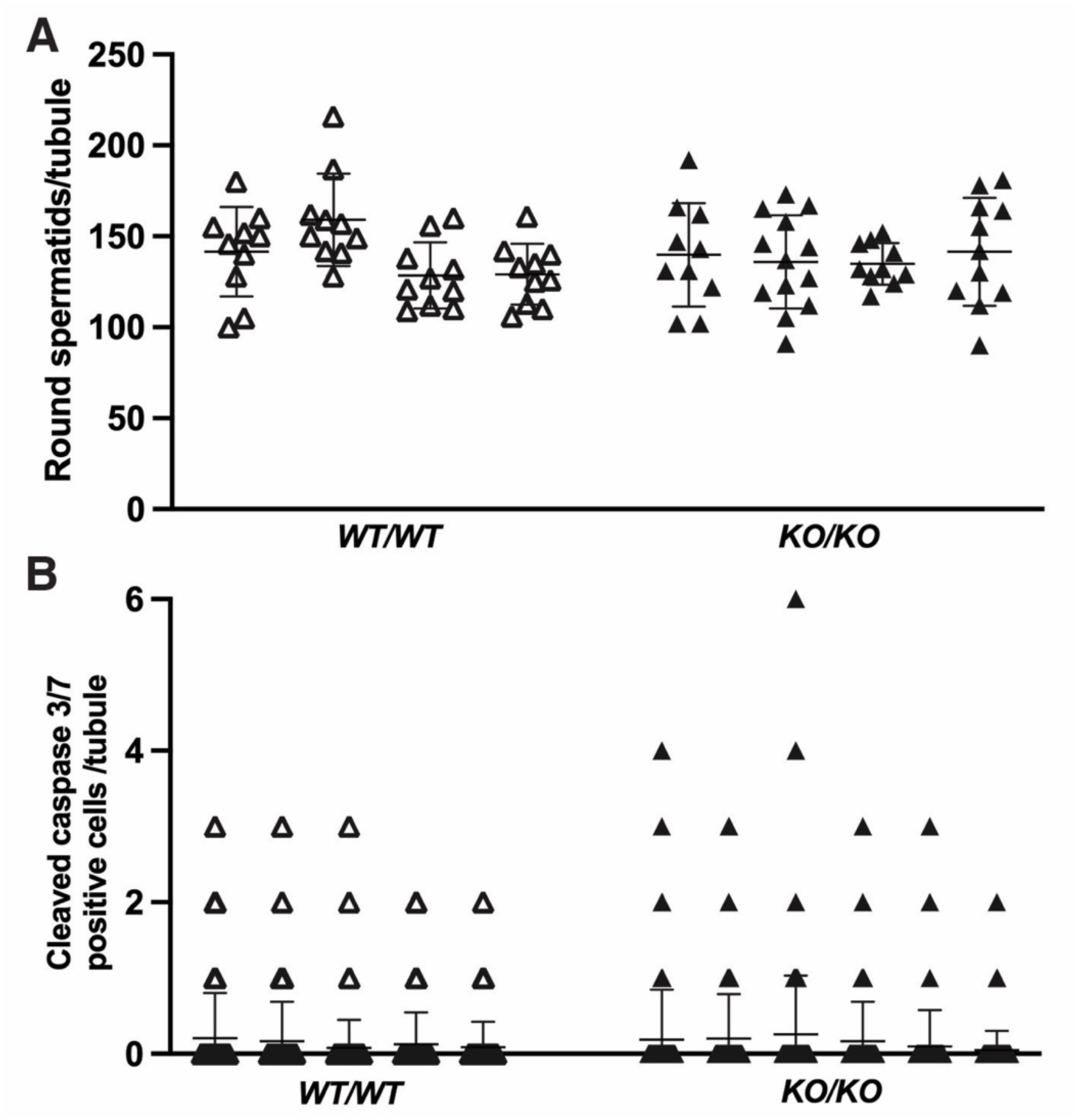
Loss of CCDC112 does not significantly affect germ cell apoptosis or round spermatid number. (A) Round spermatids number per stage VIII tubule was assessed in *Ccdc112^WT/WT^*and *Ccdc112^KO/KO^* testes (n = 4/genotype). (B) Germ cell apoptosis in *Ccdc112^KO/KO^* testes was assessed via cleaved caspase 3 and 9 staining (n = 5-6/genotype). Lines denote mean ± SD.

**Figure S3.**
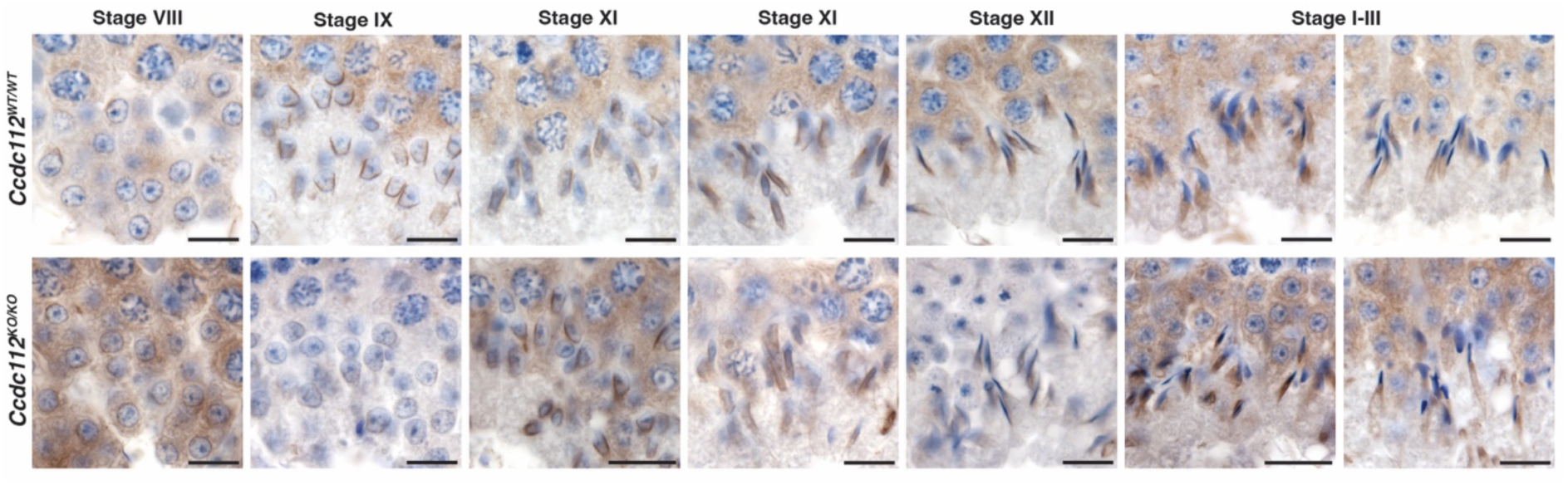
Manchette formation and movement does not require CCDC112. Assessment of manchette formation via α-tubulin, a manchette microtubule marker, staining of *Ccdc112^WT/WT^* and *Ccdc112^KO/KO^*testis sections. Manchette assembly, elongation, descension, and disassembly appears comparable between genotypes. Scale bars = 20 µm.

**Figure S4.**
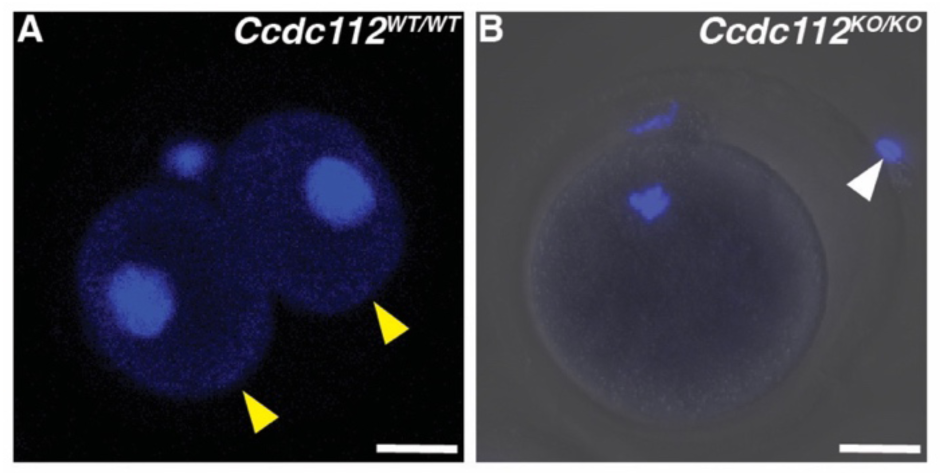
CCDC112 is essential for sperm penetration of the zonae pellucidae of oocytes *in vitro*. Immunofluorescence staining of oocytes 16 hours post-fertilisation with *Ccdc112^WT/WT^*and *Ccdc112^KO/KO^* mouse sperm as marked by DAPI. Oocytes fertilised with *Ccdc112^WT/WT^* mouse sperm developed to a two-cell stage (A; yellow arrowheads). Conversely, *Ccdc112^KO/KO^* mouse sperm were mostly unable to penetrate the zonae pellucidae (B; white arrowhead). Scale bars = 20 µm.

**Figure S5.**
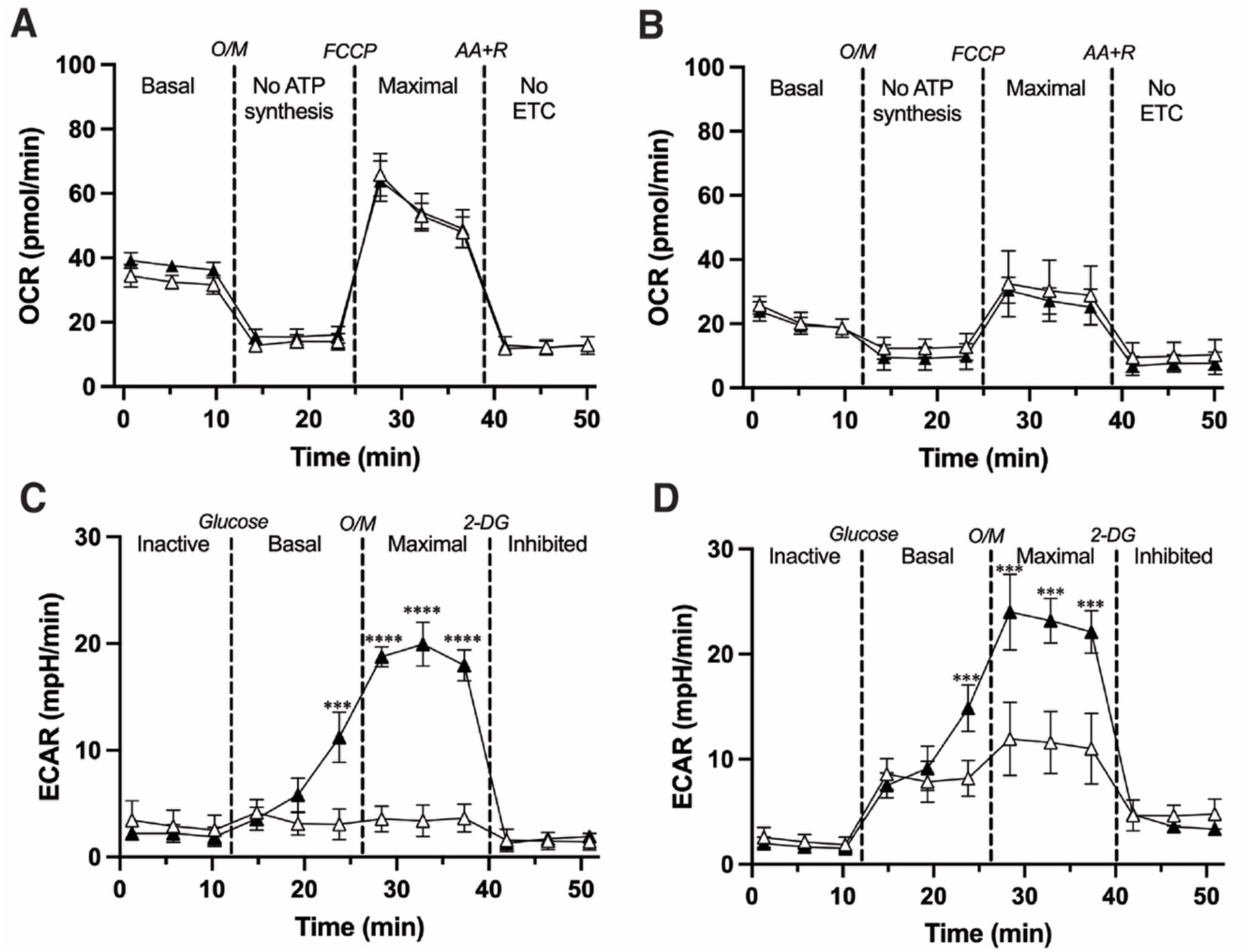
The effect of CCDC112 presence or absence in non-capacitated and capacitated sperm metabolism. Mitochondria stress test assay on *Ccdc112^WT/WT^* (A) and *Ccdc112^KO/KO^*(B) non-capacitated (white triangles) and capacitated mouse sperm (black triangles) (n = 3/genotype). Glycolytic flux Seahorse assay on *Ccdc112^WT/WT^* (C) and *Ccdc112^KO/KO^*(D) non-capacitated and capacitated mouse sperm samples (n = 3 mice/genotype). OCR = oxygen consumption rate; ECAR = extracellular acidification rate; O/M = oligomycin; AA+R = antimycin A and rotenone; 2-DG = 2-Deoxy-D-Glucose. Lines denote mean ± SD; *** P < 0.001, **** P < 0.0001.

**Supplementary videos 1 and 2. CCDC112 is essential for a normal sperm flagella waveform**

Representative videos of *Ccdc112^WT/WT^* (1) and *Ccdc112^KO/KO^* (2) mouse sperm flagella waveform over 4.5 seconds.

